# Single-molecule tweezers decoding hidden dimerization patterns of membrane proteins within lipid bilayers

**DOI:** 10.1101/2025.01.22.634249

**Authors:** Victor W. Sadongo, Eojin Kim, Seoyoon Kim, W.C. Bhashini Wijesinghe, Tae Seung Lee, Jeong-Mo Choi, Duyoung Min

## Abstract

Dimerization of transmembrane (TM) proteins is an essential biological process within cellular membranes, playing a key role in diverse pathophysiological pathways and serving as a promising therapeutic target. Although often simplified as a two-state transition from freely diffusing monomers to fully formed dimers, the dimerization process after monomer diffusion—the post-diffusion dimerization—is likely more complex due to intricate inter-residue interactions. Here, we introduce a single-molecule tweezer platform to map detailed profiles of the post-diffusion transitions in TM protein dimerization. This approach captures reversible dimerization events of a single TM dimer, revealing hidden intermediate states that emerge following the quiescent phase of monomer diffusion. Profiling the post-diffusion intermediates, kinetics, and energy landscapes—integrated with molecular dynamics simulations—uncovers the dimerization pathway, the effects of residue interactions and lipid bilayers, and the kinetic and energetic contributions of distinct dimerization domains. Furthermore, this platform characterizes selective and localized modulations via peptide binding, underscoring its potential to elucidate the mechanisms of action of TM dimer-targeting drugs at single-molecule resolution.

## Main Text

Transmembrane (TM) protein dimerization is a fundamental biological event within lipid bilayers, crucial for regulating protein activity and mediating signal transduction^1–7^. It underpins key cellular processes, including growth factor signaling via receptor tyrosine kinases, organogenesis via Notch receptors, and cell adhesion via integrins and cadherins^4,6,8–11^. Targeting TM protein dimerization offers a promising therapeutic strategy for diseases such as cancer and immune disorders^1,12–16^. For example, monoclonal antibodies have been developed to inhibit the dimerization of the epidermal growth factor receptor (EGFR) family and its downstream signaling, thereby blocking tumor growth^17–20^. Anti-TM domain peptides, such as computed helical anti-membrane proteins (CHAMP), have also been designed to inhibit the formation of TM dimers, including cytokine receptors, integrins, and G protein-coupled receptors (GPCRs)^1,12,14,15,21–25^.

Various experimental approaches, such as molecular chimera assays with dimerization reporters, fluorescence-based methods, and steric trapping, have identified interaction motifs for TM protein dimerization, characterized thermodynamic stabilities under various membrane environments, and elucidated the underlying dimerization mechanisms^2,3,5,26–40^. However, the detailed events following monomer diffusion in lipid bilayers—the post-diffusion dimerization process—remain largely unexplored (Fig. 1). The commonly assumed two-state model, in which TM monomers transition directly to fully formed TM dimers, may oversimplify the process, as computational studies suggest rugged dimerization energy landscapes with multiple intermediate states^41–46^. Furthermore, drug molecules primarily target specific residue interactions, emphasizing the need to understand the post-diffusion transition dynamics, which become even more complex in the presence of large extramembrane domains.

**Fig. 1.**
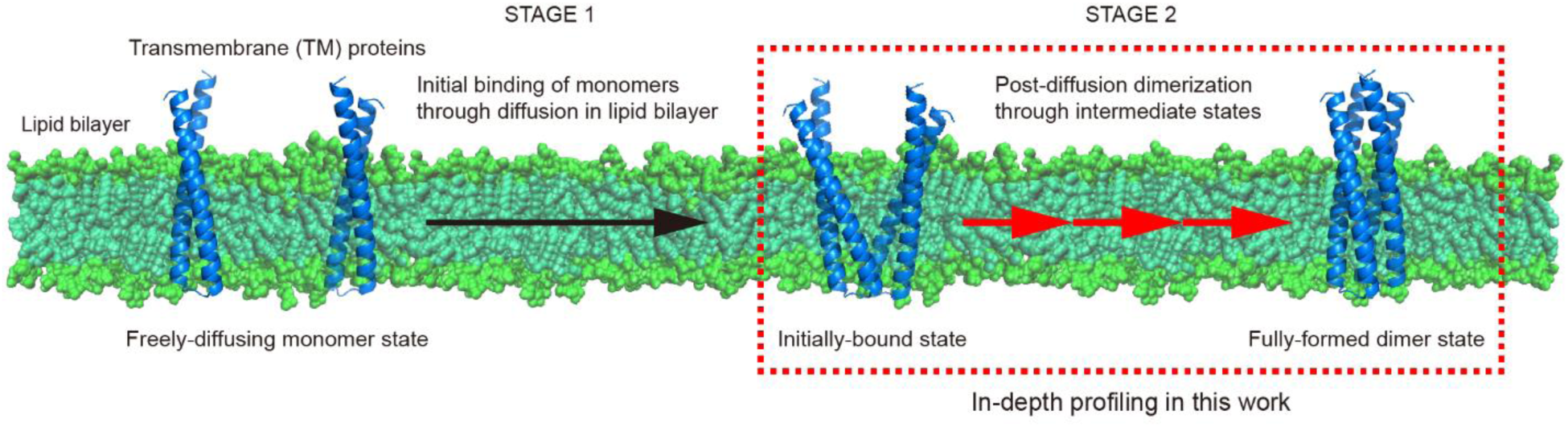
Dimerization process of transmembrane (TM) proteins. The dimerization process of transmembrane (TM) proteins can be divided into two stages: 1) the free diffusion of TM monomers within a lipid bilayer and their initial binding, and 2) the formation of dimers through intermediate states, *i.e.*, the post-diffusion dimerization. This study focuses on the second stage, highlighted by the red dashed box, and investigates it in depth using single-molecule tweezers and molecular dynamics simulations. The protein structures shown represent TMHC2, a designed TM homodimer used as a model protein in this work.

Unlike previous single-molecule force spectroscopy (SMFS) methods primarily designed to study soluble domain/protein interactions^47–61^ (see Conclusions for details), our single-molecule tweezer method presented here are tailored for TM protein interactions within lipid bilayers. Specifically, this approach profiles the post-diffusion events in TM protein dimerization by unveiling hidden intermediate states, transition kinetics, and the dimerization energy landscape after the monomer-diffusion step. Moreover, peptides targeting specific dimerization domains enable the precise examination of localized modulation patterns in TM protein dimerization. Our method offers a robust tool for unraveling the complexities of TM protein dimerization and interactions, particularly in biologically and pharmaceutically significant systems, such as receptor tyrosine kinases and cytokine receptors.

### Repetitive dissociation and dimerization of a single TM dimer within lipid bilayers

We established our single-molecule tweezer system using a designed TM homodimer, TMHC2 (Fig. 2a,b, Supplementary Figs. 1–4, and Methods). Each TMHC2 monomer consists of two TM helices forming a helical hairpin^62^ (Fig. 2a and Supplementary Fig. 4a,b). This TM dimer was chosen as a test model for two reasons: (1) experimental accessibility—its single-chained variant was recently demonstrated to be a robust model for single-molecule tweezer studies of TM protein (un)folding^63^; (2) physiological relevance—its TM domain assembly is predominantly mediated by leucine zipper-like interactions (Supplementary Fig. 4a,b), a common feature in TM dimers such as receptor tyrosine kinases, cytokine receptors, and cadherins^1,64–69^.

**Fig. 2.**
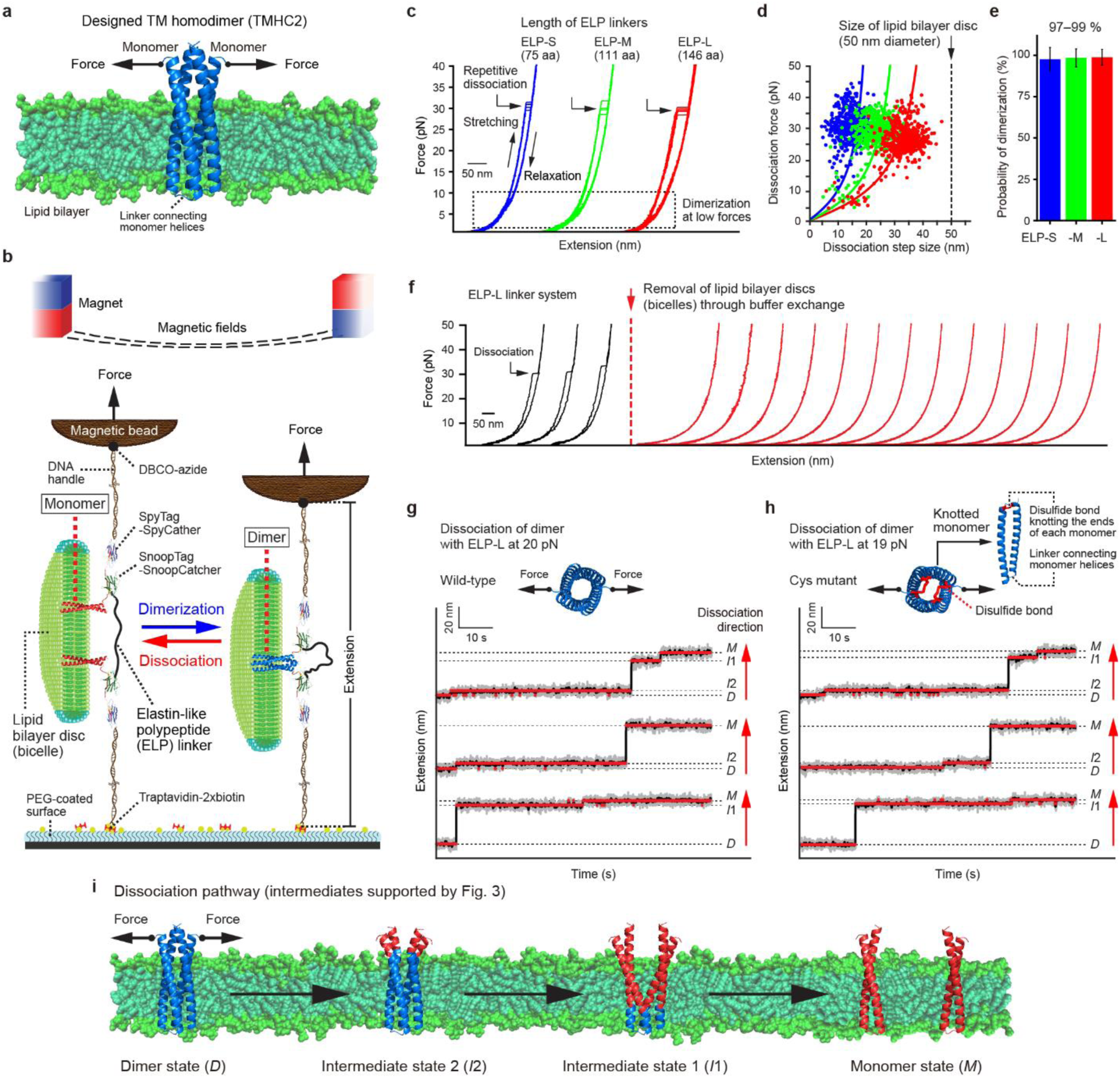
Repetitive dissociation and dimerization of a single TM dimer within lipid bilayers. (**a**) Designed transmembrane (TM) homodimer TMHC2 as a model protein. Mechanical force applied is indicated by arrows. (**b**) Our single-molecule tweezer platform. Monomers are connected by an elastin-like polypeptide (ELP) linker, enabling observation of repetitive dissociation and dimerization. (**c**) Force-extension curves showing repetitive dissociation events of a single TMHC2 during force-cycle experiments, where the force is gradually applied and released. The black dashed box highlights the force range for re-dimerization after dissociation. (**d**) Scatter plots of dissociation forces and step sizes (*n* = 500 data points from 5 molecules for each ELP linker system). The data are analyzed using the worm-like chain model (curved lines). All step sizes are smaller than the diameter of lipid bilayer discs (bicelles; 50 nm). (**e**) Probability of dimerization in the force-cycle experiments (*n* = 500 traces from 5 molecules for each ELP linker system; mean ± SD). (**f**) Force-extension curves before and after the removal of lipid bilayer discs through buffer exchange (shown in black and red, respectively). (**g**,**h**) Time-resolved extension traces showing the dissociation of wild-type TMHC2 (**g**) and its Cys mutant (**h**) during force-clamp experiments, where the force is held constant. The dimer state (*D*) transitions to the monomer state (*M*) via intermediate states (*I*_2_, *I*_1_, or both). In the Cys mutant, the N- and C-terminal ends of each monomer are knotted by a disulfide bond, shown in red. The gray, black, and red traces represent raw traces, median-filtered traces, and traces identified using a hidden Markov model, respectively. Red arrows indicate the direction of dissociation from the *D* to *M* state. (**i**) Dissociation pathway of TMHC2 observed using our single-molecule tweezers, with the intermediate positions corroborated by data shown in Fig. 3.

To apply forces to a single TM dimer, the N-terminal ends of its component monomers were respectively tethered to a glass surface and a magnetic bead via ∼1 kbp DNA handles, using orthogonal conjugation methods like DBCO-azide click chemistry, SpyCatcher-SpyTag binding, and traptavidin-2×biotin binding^63,70^ (Fig. 2b and Methods). To prevent permanent dissociation, the N-terminal ends were also connected to each other with an elastin-like polypeptide (ELP) linker that lacks secondary structures, via SnoopCatcher-SnoopTag binding (Fig. 2b, Supplementary Figs. 1–3, and Methods). The ELP linker enables repetitive dimerization after force-induced dissociation (Fig. 2c; see below for details). We used ELP linkers of varying lengths: 75, 111, and 146 aa (ELP-S, -M, and -L; Supplementary Fig. 1). The SnoopCatcher (with a disulfide knot positioned appropriately) and SpyCatcher proteins are covalently complexed with their partner peptide tags, remaining folded under applied forces^63,70,71^ (Supplementary Fig. 3). The lipid bilayer environment was reconstituted using bicelles, lipid bilayer discs surrounded by detergents (Fig. 2b and Methods).

Gradual force application dissociated single TM dimers into monomers at ∼20–40 pN, as indicated by stepwise increases in molecular extension due to ELP linker stretching (Fig. 2c). As expected, the dissociation step size (∼10–40 nm) increased with ELP linker length, consistent with the predictions of the worm-like chain model^72^ (Fig. 2c,d, Supplementary Fig. 5, and Methods). Gradual force relaxation back to 1 pN induced dimer reformation (Fig. 2c), achieving 97–99% efficiency in repetitive force cycles (Fig. 2e). In these force-cycle experiments, dimerization occurred at low force ranges (<10 pN), where step sizes shortened below ∼20 nm (Fig. 2c,d). Across all ELP linkers of varying lengths, the 10–40 nm dissociation/dimerization step sizes fell within the diameter of lipid bilayer discs (∼50 nm) (Fig. 2d and Supplementary Figs. 5 and 6), suggesting that these events occurred in a lipid bilayer. Indeed, dissociation events were not detected following the removal of lipid bilayer discs through buffer exchange, indicating a failure of re-dimerization in the absence of lipid bilayers (Fig. 2f and Supplementary Fig. 7).

### Identifying hidden intermediate states in TM protein dimerization

In the force-cycle experiments, potential dissociation intermediates may have been obscured by rapid transitions resulting from high force levels. We thus turned to force-clamp experiments at constant lower forces, slowing transitions and revealing previously hidden intermediates (Fig. 2g,h). At 20 pN, we identified one or both of two intermediates during dimer dissociation (Fig. 2g and Supplementary Fig. 8a). A Cys mutant, containing a disulfide bond that links the N- and C-terminal ends of each monomer, exhibited a similar dissociation pattern (Fig. 2h and Supplementary Fig. 8b). This observation suggests that the intermediates arise from interactions between monomers rather than intra-chain interactions between helices within each monomer, which aligns with our pulling design that does not separate the monomer helices. The pulling geometry also dictates a specific dissociation pathway, progressing from the pulling side to the opposite side (Fig. 2i; intermediate positions supported by Fig. 3).

**Fig. 3.**
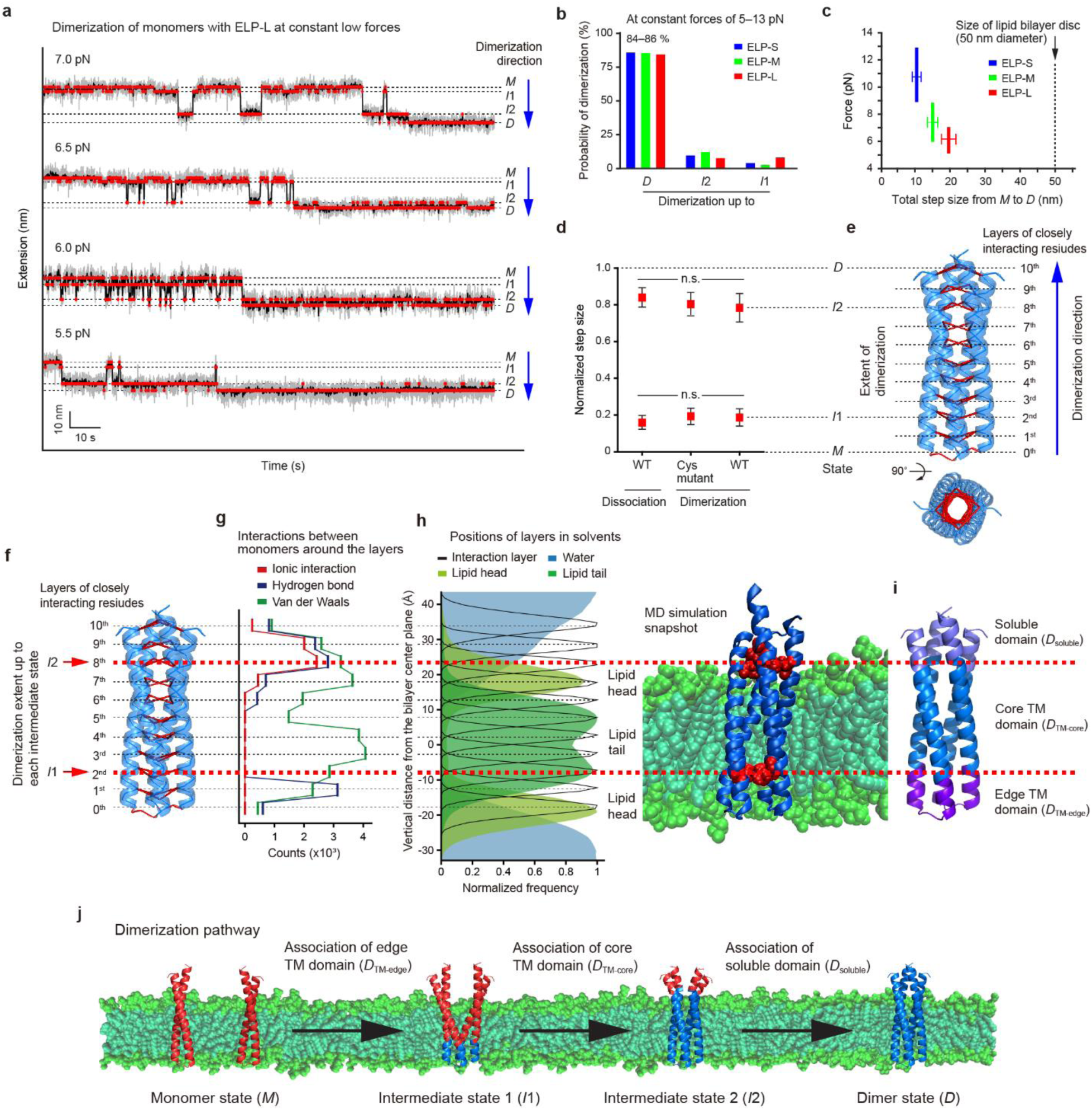
Hidden intermediate states in TM protein dimerization. (**a**) Time-resolved extension traces showing dimerization events of a single TMHC2 during force-clamp experiments, where the force is held constant. The monomer state (*M*) transitions to the dimer state (*D*) states via two intermediate states (*I*_1_ and *I*_2_). The gray, black, and red traces represent raw traces, median-filtered traces, and traces identified using a hidden Markov model, respectively. Blue arrows indicate the direction of dimerization from the *M* to *D* state. (**b**) Probability of dimerization up to each state in the force-clamp experiments (*n* = 224–358 traces from 8–13 molecules). (**c**) Distribution of total step sizes from the *M* to *D* state (*n* = 142–238 data points from 8–13 molecules; mean ± SD). All step sizes are smaller than the diameter of lipid bilayer discs (bicelles; 50 nm). (**d**) Normalized step sizes for the intermediates. The data are presented as mean ± SD (right, *n* = 530 data points from 34 molecules in all ELP systems at 5–13 pN; middle, *n* = 59 data points from 3 molecules in the ELP-L system at 5–7 pN; left, *n* = 7–9 data points from 3 molecules in the ELP-L system at 20 pN). One-way ANOVA with post-hoc Tukey HSD test (n.s. for *p* > 0.05). (**e**,**f**) TMHC2 dimer structure showing the layers of closely interacting inner residues at the dimerization interface. The normalized step sizes of the intermediates in panel **d** are compared to the extent of dimerization from the 0^th^ to 10^th^ layer in panel **e**. (**g**) Counts of inter-monomer interactions around the interaction layers. This data was obtained using RING and all-atom molecular dynamics (AAMD) simulations. (**h**) Vertical positions of the interaction layers compared to solvent components. The data on the left was obtained from AAMD simulations, with a simulation snapshot shown on the right. (**i**) Dimerization domains separated by the intermediate positions. (**j**) Dimerization pathway of TMHC2 observed using our single-molecule tweezers.

To investigate intermediates during dimerization, we reduced the force to 5–13 pN following complete dissociation at 35 pN (Fig. 3a and Supplementary Fig. 9). At these low forces, transitions between different extension levels were frequent but not clearly resolved due to high fluctuation noise from the tethered bead (Fig. 3a and Supplementary Figs. 10–13). To address this, we utilized the Bayesian information criterion (BIC) and hidden Markov modeling (HMM), thereby identifying four distinct states: monomer (*M*), intermediate 1 (*I*_1_), intermediate 2 (*I*_2_), and dimer (*D*) (Fig. 3a, Supplementary Figs. 14–16, and Methods). Even at forces of 5–13 pN, 84–86% of traces showed full dimerization to the *D* state, strictly following the sequence *M*→*I*_1_→*I*_2_→*D* with no bypasses (Fig. 3b, Supplementary Fig. 17, and Supplementary Table 1). The total dimerization step sizes (∼10–20 nm) fell within the diameter of lipid bilayer discs (∼50 nm) (Fig. 3c and Supplementary Figs. 15 and 16), consistent with the in-bilayer transitions discussed above.

The pulling geometry restricts dimerization to a specific pathway, progressing from the opposite side to the pulling side—*i.e.*, the reverse of dissociation (Fig. 2i). However, our pulling design captures the predominant dimerization pathway in the absence of external forces (see subsequent sections for details). When the extension levels for the *M* and *D* states were normalized to 0 and 1, respectively, the *I*_1_ and *I*_2_ states corresponded to ∼0.2 and ∼0.8 on average (Fig. 3d and Supplementary Fig. 18). These intermediate positions closely matched those observed during wild-type dissociation and Cys mutant dimerization (Fig. 3d and Supplementary Fig. 18). These results indicate that the intermediates *I*_1_ and *I*_2_ arise from monomer interactions, representing ∼20% and ∼80% dimer formation.

### Structural mapping of dimerization intermediate states

To investigate the structures of the intermediate states (Fig. 3e–j and Methods), we designated the leucine zipper-like, closely interacting inner residues at the dimerization interface as the 1^st^ to 10^th^ interaction layers, spanning every 3–4 residues from the opposite side to the pulling side (Fig. 3e,f and Supplementary Fig. 4a,b). The polypeptide linkers connecting the monomer helices were also designated as the 0^th^ interaction layer due to their flexibility, allowing close interactions. Based on the dimerization direction (the 0^th^ to 10^th^ layer), the *I*_1_ and *I*_2_ states with ∼20% and ∼80% dimer formation approximately correspond to dimerization up to the 2^nd^ and 8^th^ layers, respectively (Fig. 3e,f and Supplementary Fig. 4a,b). This structural mapping of intermediates, combined with all-atom molecular dynamics (AAMD) simulations, identifies three distinct domains: edge TM domain (*D*_TM-edge_, 0^th^–2^nd^ layers), core TM domain (*D*_TM-core_, 2^nd^–8^th^ layers), and soluble domain (*D*_soluble_, 8^th^–10^th^ layers) (Fig. 3h,i and Methods). The dimerization pathway thus follows sequential association of the *D*_TM-edge_, *D*_TM-core_, and *D*_soluble_ domains (Fig. 3j).

Using AAMD simulations and the residue interaction network generator (RING)^70,73,74^ (Methods), we analyzed factors determining intermediate positions in the inter-monomer interactions (Fig. 3g,h). Van der Waals interactions (*F*_vdw_) are broadly distributed across the layers, with peaks at the 3^rd^, 4^th^, and 7^th^ layers (Fig. 3g and Supplementary Fig. 19). Other intermolecular forces dominate only in specific regions: hydrogen bonding (*F*_H-bond_) in the 0^th^–1^st^ and 8^th^–9^th^ layer regions, and ionic interactions (*F*_ionic_) in the 8^th^–9^th^ layer regions. This AAMD-RING analysis suggests that intermediate positions occur in regions with significant changes in specific residue interactions—*i.e.*, the transition from *F*_H-bond_ to *F*_vdw_ at the ∼2^nd^ layer and the sharp emergence of *F*_H-bond_ and *F*_ionic_ at the ∼8^th^ layer (Fig. 3g). In contrast, the intermediate positions are not strictly associated with solvent boundaries: the 8^th^ layer lies at the lipid head-water interface, whereas the 2^nd^ layer is embedded within the lipid tail region (Fig. 3h).

### Our method observes predominant dimerization pathway under force-free conditions

The natural dimerization process, in the absence of external forces, is not necessarily restricted to a specific pathway, such as the 0^th^-to-10^th^ layer dimerization discussed above. To see whether our method observes the predominant pathway under force-free conditions, we conducted coarse-grained MD (CGMD) simulations using the Martini model for two initial systems (Fig. 4a and Methods): (1) initial monomer system, with unbound monomers separated by ∼10 nm in a lipid bilayer, allowing free diffusion and dimer formation, and (2) initial dimer system as a control. To confirm stable dimer formation in the monomer system, we calculated the root mean square deviation (RMSD) from the final protein structure at 2 μs for both systems (Fig. 4b). After dimer formation (see Methods for detailed criterion), the RMSDs were comparable between the two systems, indicating that monomers stably formed dimers (Fig. 4c).

**Fig. 4.**
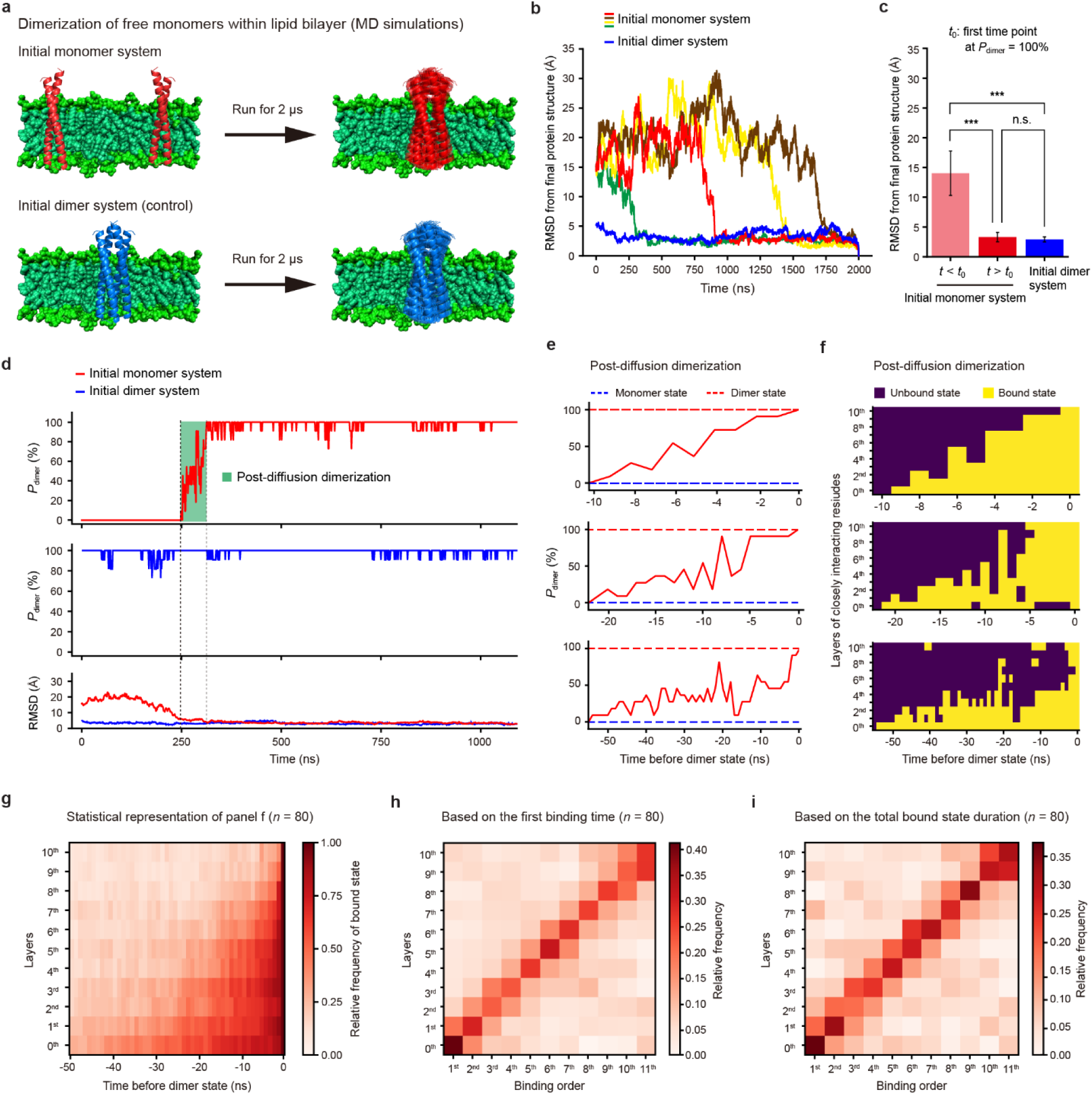
Predominant dimerization pathway under force-free conditions. (**a**) Coarse-grained molecular dynamics (CGMD) simulations to identify the predominant dimerization pathway of TMHC2. (**b**) Root mean square deviation (RMSD) from the final protein structure at 2 μs. (**c**) Average RMSD for the initial monomer system (*n* = 80 simulations; mean ± SD) and the initial dimer systems (*n* = 5 simulations; mean ± SD). One-way ANOVA with post-hoc Tukey HSD test (*** for *p* < 0.001 and n.s. for *p* > 0.05). *P*_dimer_ indicates the percentage of dimerization. (**d**) Percentage of dimerization (*P*_dimer_) as a function of time. Representative *P*_dimer_ traces are shown for both the initial monomer and dimer systems, with RMSD shown at the bottom for comparison. (**e**) Percentage of dimerization (*P*_dimer_) during the post-diffusion dimerization period (green-boxed region in panel **d**). Three representative traces are shown. (**f**) Barcode plots for binding at each interaction layer. The plots are derived from the same trajectories analyzed in panel **e**. (**g**) Statistical representation for binding at each interaction layer during the post-diffusion dimerization period (*n* = 80 simulations). Values of 0 and 1 are assigned to the unbound and bound regions in panel **f**, respectively, and then averaged across 80 trajectories. (**h,i**) Binding order across the interaction layers (*n* = 80 simulations). The binding order is analyzed using two different criteria: the first binding time and the total bound state duration during the post-diffusion dimerization period.

We then quantified the dimerization process after monomer diffusion—*i.e.*, the post-diffusion dimerization (Methods). Following monomer diffusion, the dimerization percentage increased rapidly from 0% to 100% (Fig. 4d,e). During this phase, time-resolved binding profiles were characterized for the 0^th^–10^th^ interaction layers (Fig. 4f and Supplementary Fig. 20). Across 80 simulations showing dimer formation, a statistically predominant binding order was observed, proceeding sequentially from the 0^th^ to 10^th^ layer (Fig. 4g). This binding order was further validated through quantitative analyses of the first binding time or total bound state duration (Fig. 4h,i and Supplementary Fig. 21). This directional binding is likely driven by the asymmetric solvation of hydrophilic interaction layers within the hydrophobic lipid membrane—specifically, the 0^th^–1^st^ layers likely serve as nucleation points due to their strong hydrogen bonding within the lipid membrane (Fig. 3f–h). These results suggest that our tweezer method captures the transitions along the predominant dimerization pathway (the 0^th^ to 10^th^ layer) in the absence of external forces.

### Mapping post-diffusion kinetics and energy landscape in TM protein dimerization

We revisited the single-molecule experimental data to estimate the zero-force, post-diffusion kinetics and energetics of the dimerization transitions (Fig. 5, Supplementary Table 2, and Methods). Using the Bell-Evans model^72^, the force-dependent rate constants for each dimerization domain (*k*_a_-association, *k*_d_-dissociation) were extrapolated to obtain the zero-force rate constants (*k*_0_) and the distances to the transition states (Δ*x*^†^) (Fig. 5a and Supplementary Fig. 22). By further correcting for ELP linker length to the minimum monomer distance (Supplementary Fig. 23), we estimated the post-diffusion transition kinetics at zero force (*k*_0,m_, Δ*x*^†^_m_) (Fig. 5b).

**Fig. 5.**
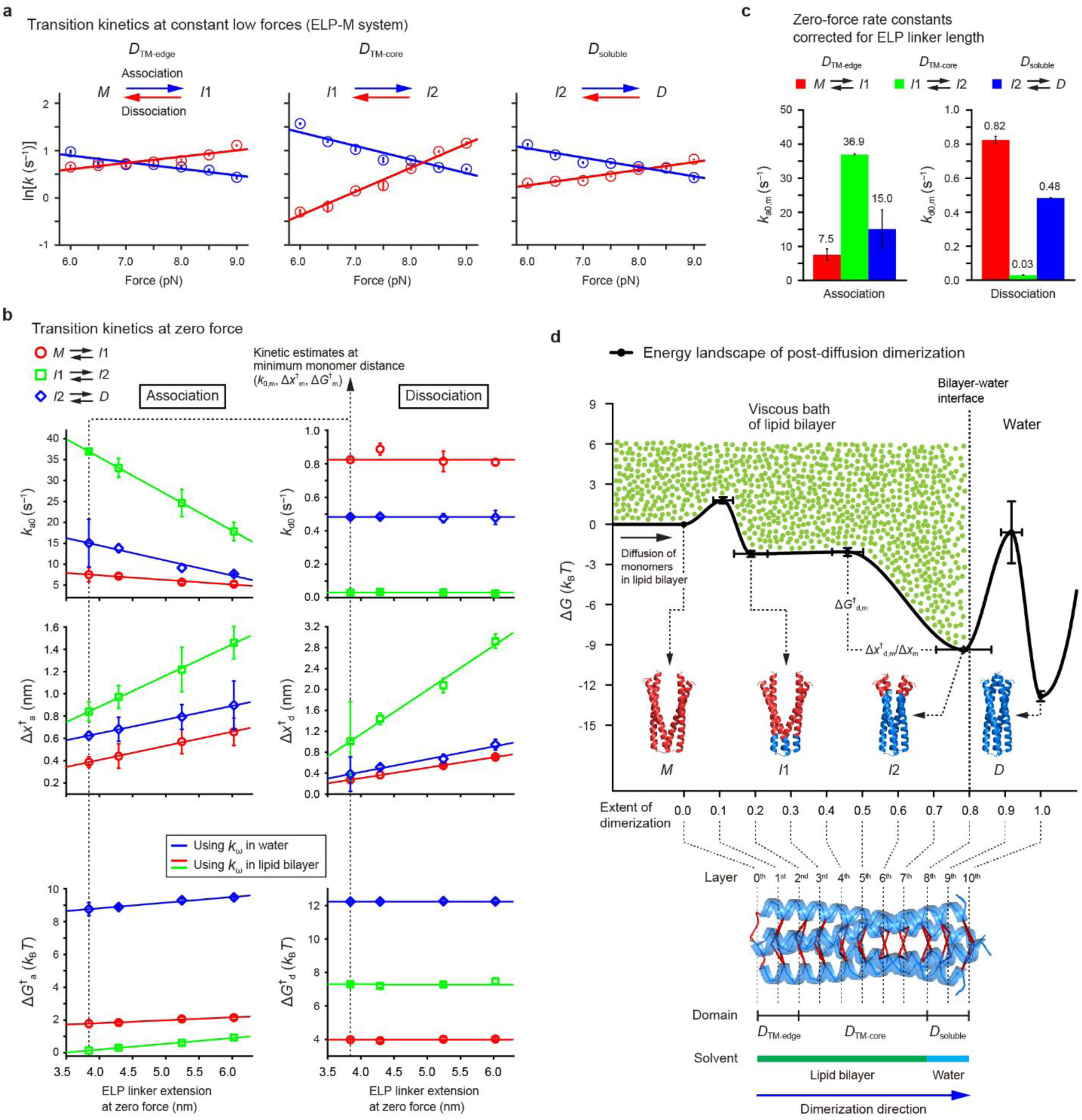
Post-diffusion kinetics and energy landscape in TM protein dimerization. (**a**) Rate constants (*k*) for transitions in TMHC2 as a function of force. The data correspond to those in the ELP-M linker system, with association and dissociation indicated in blue and red, respectively. The symbols *M*, *I*_1_, *I*_2_, and *D* indicate the monomer, intermediate 1, intermediate 2, and dimer states, respectively. The error bars indicate SE. (**b**) Transition kinetics at zero force as a function of ELP linker extension. The zero-force kinetics (*k*_0_ and 𝛥*x*^†^) are obtained from extrapolation analysis of the data in panel **a**, where *k*_0_ and 𝛥*x*^†^ are the rate constant at zero force and the distance to the transition state, respectively. The energy barrier height (𝛥*G*^†^) is derived from *k*_0_ using the Kramers rate framework, with different values of the frequency factor (*k*_ω_) depending on whether the transitions occur in water or lipid bilayers. The data in this panel are extrapolated to the minimum distance between monomers to obtain the zero-force transition kinetics with ELP linker length corrected (*k*_0,m_, 𝛥*x*^†^_m_, and 𝛥*G*^†^_m_). The error bars indicate SE. (**c**) Zero-force rate constant corrected for ELP linker length (*k*_0,m_). The error bars indicate SE. (**d**) Reconstructed free energy landscape (𝛥*G*) of post-diffusion TMHC2 dimerization. The reaction coordinate represents the extent of dimerization from the 0^th^ to 10^th^ interaction layer. The green dots illustrate the high viscosity of the lipid bilayer. The bilayer-water interface is denoted by a vertical dashed line. The TMHC2 dimer structure and designated interaction layers shown at the bottom serve as the structural reference for the extent of dimerization. The regions of dimerization domains and corresponding solvents are indicated below the protein structure. The symbols *D*_TM-edge_, *D*_TM-core_, and *D*_soluble_ represent the respective dimerization domains: the edge transmembrane (TM) domain, core TM domain, and soluble domain. The blue arrow indicates the direction of dimerization from the *M* to *D* state.

The initial *D*_TM-edge_ association (*M*→*I*_1_) is rate-limiting (*k*_a0,m_ = 7.5 s^−1^), likely due to monomer binding initiation (Fig. 5c). Subsequent associations are faster: *k*_a0,m_ = 36.9 s^−1^ for the *D*_TM-core_ domain (*I*_1_→*I*_2_) and *k*_a0,m_ = 15.0 s^−1^ for the *D*_soluble_ domain (*I*_2_→*D*). Dissociations are much slower than associations across all domains (*k*_d0,m_’s < 1 s^−1^), indicating that the domains are kinetically stable. The *D*_TM-core_ domain dissociates particularly slowly (*k*_d0,m_ = 0.03 s^−1^), ∼16–27 times slower than other domains, making it rate-limiting for dissociation. Notably, complete dimer dissociation (*D*→*M*) is ∼10³ times slower than the *D*_TM-core_ dissociation (Supplementary Fig. 9). The high kinetic stability of the entire dimer likely arises from the rapid re-association of domains before complete dissociation, particularly for the *D*_TM-core_ domain, which re-associates ∼10³times faster than it dissociates (Fig. 5c).

Based on the kinetic estimates and intermediate positions, we reconstructed the free energy landscape for the post-diffusion dimerization (Fig. 5d and Supplementary Fig. 24). Using the Kramers rate framework^75^, the post-diffusion rate constants (*k*_0,m_’s) were converted to energy barrier heights (Δ*G*^†^_m_’s): *k*_0,m_ = *k*_ω_·exp(–Δ*G*^†^_m_/*k*_B_*T*), where *k*_ω_ is the frequency factor. Different *k*_ω_ values were used to reflect solvent viscosity^63,76^—*k*_ω_ = 10^4^–10^6^ s^−1^ for soluble domains in water^77^ and *k*_ω_ = ∼45 s^−1^ for TM domains in lipid bilayers^63^. Transition state positions were estimated from Δ*x*^†^_m_’s (Methods). The initial *D*_TM-edge_ association (*M*→*I*_1_) requires minimal energy (Δ*G*^†^_a,m_ = 1.8 *k*_B_*T*), slightly exceeding thermal fluctuations (∼1 *k*_B_*T*). The next *D*_TM-core_ association (*I*_1_ → *I*_2_) proceeds ‘downhill’ with a barrier of only 0.1 *k*_B_*T*, and once bound, it contributes significantly to the dimer’s thermodynamic stability (Δ*G*_m,TM-core_ = 7.2 *k*_B_*T vs* Δ*G*_m,others_ = 2.3–3.5 *k*_B_*T*). The final *D*_soluble_ association (*I*_2_→*D*) requires a substantial energy of 8.8 *k*_B_*T*, completing the dimerization. Once fully dimerized, the dimer achieves high kinetic and thermodynamic stability (Δ*G*^†^_d,m,D-I2_ = 12.2 *k*_B_*T* and Δ*G*_m,D-M_ = 12.8 *k*_B_*T*), with the highest dissociation barrier near the dimer state, ensuring its structural rigidity.

Ostensibly, the barrier height estimates contradict the rate constant estimates—while the *D*_TM-edge_ domain is rate-limiting in associations (Fig. 5c), the *D*_soluble_ domain is energetically limiting (Fig. 5d). However, this apparent mismatch arises from differences in solvent viscosity between water and lipid bilayers, as reflected in the frequency factor (*k*_ω_). The high viscosity of lipid bilayers slows molecular transitions significantly, by a factor of ∼10^2^–10^4^ compared to water^63,76^, making a TM domain rate-liming. Similarly, the *D*_TM-core_ domain is rate-limiting in dissociations (Fig. 5c), while the *D*_soluble_ domain is energetically limiting (Fig. 5d). Compared to the TM domains (*D*_TM-edge_ and *D*_TM-core_), the high energy barrier for the *D*_soluble_ association may result from large thermal fluctuations in water, hindering native residue contacts. Conversely, its high dissociation barrier likely arises from strong native interactions, including electrostatic forces, hydrogen bonding, and van der Waals interactions (Fig. 3g).

### Profiling peptide-mediated modulation of TM protein dimerization

Drug molecules primarily target the post-diffusion dimerization events through intermolecular interactions. We thus assessed our method’s ability to detect drug-induced subtle changes using short extraneous peptides (Fig. 6a and Supplementary Fig. 4c): 7-aa peptide I, which binds to the 9^th^–10^th^ layers to selectively target the *D*_soluble_ domain, and 13-aa peptide II, which binds to the 7^th^– 10^th^ layers and slightly extend into the *D*_TM-core_ domain, covering only ∼23% of it. These peptides are expected to modulate downstream transitions without complete inhibition, thereby inducing localized effects (Fig. 6a). In the force-cycle experiments with 10 μM peptide (Fig. 6b), peptide I marginally reduced the probability of complete dimerization to ∼93%, whereas peptide II significantly reduced it to ∼38% (Fig. 6c and Methods). For peptide II, the remaining ∼62% exhibited partial dimerization, characterized by aberrant dissociation patterns with either small or undetectable step sizes (Fig. 6d and Supplementary Fig. 25).

**Fig. 6.**
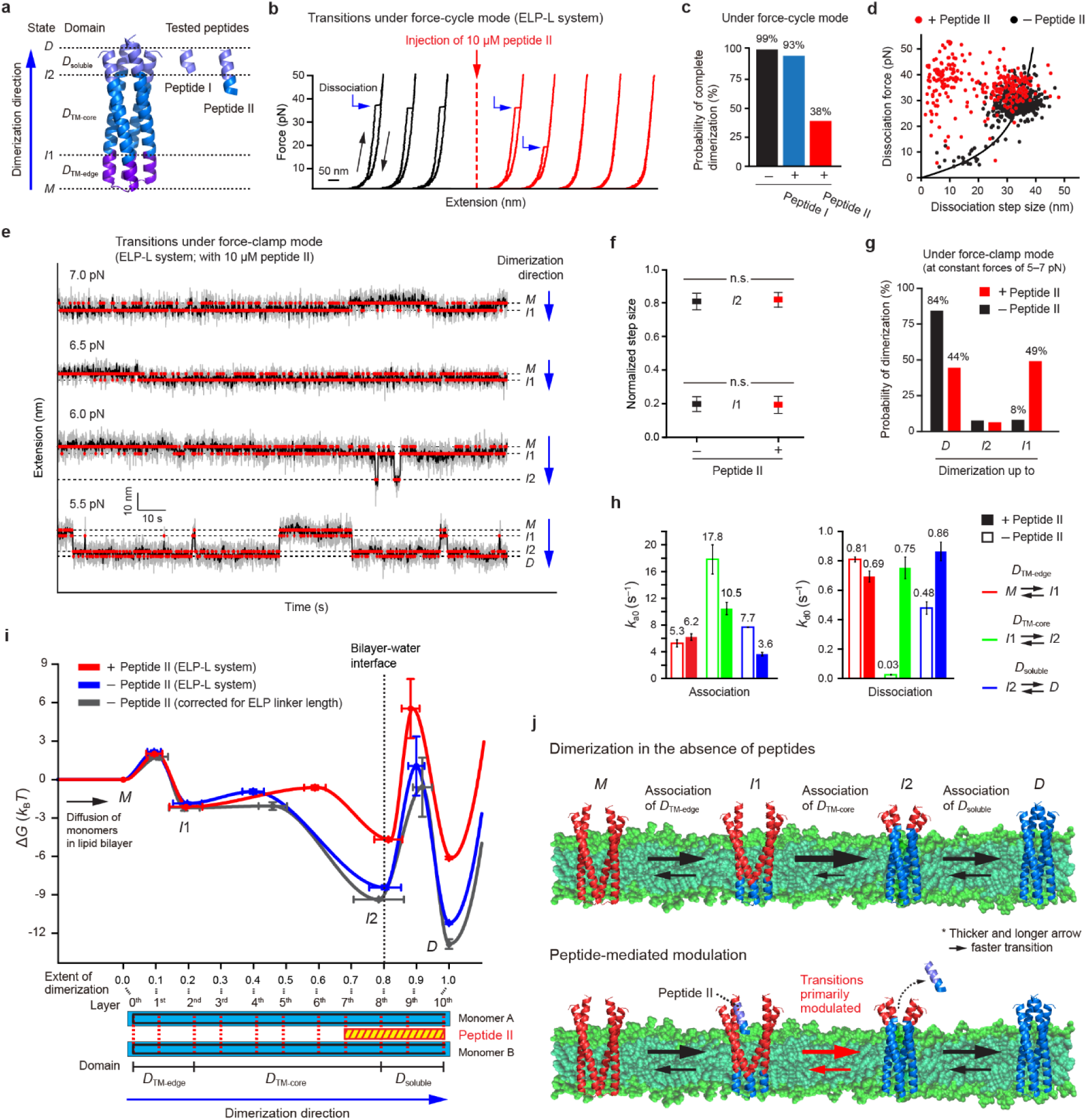
Peptide-mediated modulation of TM protein dimerization. (**a**) TMHC2 dimer and tested peptides (peptide I and II). The symbols *D*_soluble_, *D*_TM-core_, and *D*_TM-edge_ represent the respective dimerization domains: the soluble domain, core transmembrane (TM) domain, and edge TM domain. The domain regions targeted by the peptides are illustrated in different colors. The symbols *D*, *I*_2_, *I*_1_, and *M* indicate the dimer, intermediate 2, intermediate 1, and monomer states, respectively. (**b**) Force-extension curves of a single TMHC2 before and after the injection of peptide II. (**c**) Probability of complete dimerization before and after peptide injection in the force-cycle experiments (*n* = 432–513 traces from 5–7 molecules). (**d**) Scatter plots of dissociation forces and step sizes in the presence or absence of peptide II (*n* = 432–500 data points from 5–7 molecules). The data are analyzed using the worm-like chain model (curved line). (**e**) Time-resolved extension traces of a single TMHC2 in the presence of peptide II. The gray, black, and red traces represent raw traces, median-filtered traces, and traces identified using a hidden Markov model, respectively. (**f**) Normalized step sizes for the intermediates at 5–7 pN (mean ± SD; *n* = 107–142 data points from 12–13 molecules). One-way ANOVA with post-hoc Tukey HSD test (n.s. for *p* > 0.05). (**g**) Probability of dimerization up to each state in the force-clamp experiments (*n* = 224–342 traces from 12–13 molecules). (**h**) Zero-force rate constants (*k*_0_) in the presence or absence of peptide II under the ELP-L linker system. The error bars indicate SE. (**i**) Reconstructed free energy landscape (𝛥*G*) of TMHC2 dimerization in the presence or absence of peptide II. The reaction coordinate represents the extent of dimerization from the 0^th^ to 10^th^ layer. The TMHC2 dimer structure is shown at the bottom, with the interaction layers and the binding position of peptide II indicated. (**j**) Peptide-mediated modulation of TMHC2 dimerization at the level of dimerization domains. Thicker and longer arrows represent faster transitions.

We further conducted an in-depth profiling of peptide II-mediated changes in the intermediates, transition kinetics, and free energies using the force-clamp experiments (Fig. 6e, Supplementary Figs. 14, 16, and 26, and Supplementary Table 2). Although peptide II did not alter the intermediate positions (Fig. 6f and Supplementary Fig. 18), it significantly enhanced partial dimerization up to the *I*_1_ state by 41% (Fig. 6g and Supplementary Fig. 17). These results suggest that peptide II primarily traps the dimerization process in the partially formed *I*_1_ state, corresponding only to the initial *D*_TM-edge_ association. Moreover, while the transition kinetics of the *D*_TM-edge_ domain remained largely unaffected (Fig. 6h and Supplementary Fig. 22), the peptide delayed the subsequent associations of the *D*_TM-core_ and *D*_soluble_ domains by ∼2-fold and facilitated the dissociation of the *D*_soluble_ domain to a similar extent. Notably, the dissociation of the *D*_TM-core_ domain was accelerated ∼25-fold (Fig. 6h), indicating that the peptide predominantly disrupts the *D*_TM-core_ domain by greatly increasing its dissociation rate.

Peptide II significantly altered the dimerization energy landscape, primarily targeting the *D*_TM-core_ domain (Fig. 6i). While it had no effect on the *I*_1_ state (ΔΔ*G*_M-I1_ = –0.2 *k*_B_*T*), it markedly destabilized the *I*_2_ state (ΔΔ*G*_I1-I2_ = +4.0 *k*_B_*T*) by reducing the dissociation barrier height of the *D*_TM-core_ domain by 3.4 *k*_B_*T*. Additionally, the peptide shifted the transition state position of the *D*_TM-core_ domain from the 4^th^ to the 6^th^ layer (Fig. 6i). This transition state shift aligns with the requirement for an extended association up to the 6^th^ layer to displace the bound peptide at the 7^th^– 10^th^ layers (Fig. 6i, inset). The energy landscape for the final *D*_soluble_ domain shifted upward while retaining its overall shape, reflecting solely the propagation of earlier changes in the *D*_TM-core_ domain. Thus, peptide II selectively disrupted the *D*_TM-core_ domain in both kinetic and thermodynamic dimensions (Fig. 6i,j). Despite binding to only 23% of the domain, it caused a dramatic inhibition of complete dimerization.

## Conclusions

Various SMFS methods, based on magnetic/optical tweezers or atomic force microscopy (AFM), have been developed to investigate protein-protein interactions at single-molecule resolution. These approaches are primarily limited to interactions involving water-soluble proteins/peptides or the soluble regions of TM proteins^47–61^. In contrast, our tweezer approach enables precise tracking of TM protein interactions within lipid bilayers with unprecedented detail. Additionally, it offers a modular single-molecule tweezer platform applicable to studying a wide range of protein-protein interactions. In the systems such as magnetic/optical tweezers, tethering target molecules with linkers, such as polypeptide linkers, is essential to prevent permanent dissociation and allow repetitive association cycles. A convenient strategy is to genetically fuse proteins with a polypeptide linker^47,50,52,53,61^. However, this approach often results in low expression levels, particularly for TM proteins, making fused constructs challenging to produce. To address this issue, our method employs an ELP linker flanked by SnoopCatchers, which is easily expressed and purified in large quantities. The SnoopCatcher proteins then rapidly and covalently bind to SnoopTags at the ends of target TM proteins.

TM protein dimerization is often modeled as a two-state process, where monomers diffuse and transition into dimers in one simple step. However, this model oversimplifies the process, as monomer diffusion does not determine detailed association patterns following the monomers’ initial binding. Understanding these residue-level interactions is crucial, as drugs primarily target the post-diffusion binding events rather than the diffusion process itself. Our tweezer platform enables detailed mapping of the post-diffusion transitions within lipid bilayers, offering structural, kinetic, and thermodynamic insights into specific dimerization domains. Even with a simplified TM dimer model, our approach reveals broad insights that are generally applicable: 1) dimerization domains are primarily dictated by changes in specific residue interactions rather than solely by bilayer-water interfaces; 2) TM domains are more likely to be rate-limiting in dimerization due to the high viscosity of lipid bilayers, while water-soluble domains are more likely to be energetically limiting due to the high thermal fluctuations in water; 3) the local kinetic and energetic contributions of dimerization domains can vary, providing opportunities for precise modulation of dimerization through molecular agents.

Our method is broadly applicable to complex TM protein dimerization and interaction systems of biological and pharmaceutical significance. For example, the hidden dynamics of cancer-associated, ligand-induced EGFR family dimerization could be elucidated using our platform. With the growing recognition of designed peptides as therapeutic tools for inhibiting TM dimerization^1,12,14,15,21–25^, our approach can capture subtle, peptide-mediated modulation at the level of dimerization domains. For instance, the soluble portion of a tested peptide, characterized by high aqueous solubility, acts as a binding anchor to the target’s soluble domain, while its minimal TM segment disrupts both the kinetic and thermodynamic stability of the target’s TM domain, significantly inhibiting complete dimerization. Our tweezer approach thus offers a powerful tool for unraveling the mechanisms of action of TM dimer-targeting drugs at single-molecule resolution.

## Methods

### Protein expression and purification

The monomeric construct of the designed transmembrane (TM) homodimer TMHC2^62^ studied in this work contains SpyTag and SnoopTag sequences at its N-terminus^78,79^, linked by GGSGGS linkers (Supplementary Fig. 1). These peptide tags enable the attachment of DNA handles via SpyTag-SpyCatcher binding^63,78,80^, and the attachment of elastin-like polypeptide (ELP) linkers via SnoopTag-SnoopCatcher binding^59,79^ (Supplementary Fig. 3). For the ELP linker constructs, the ELP sequences of three different lengths were inserted between two SnoopCatcher sequences (Supplementary Fig. 1). To prevent unwanted unfolding of the C-terminal SnoopCatcher, cysteine residues were introduced at the start and end positions of its sequence, enabling the spontaneous formation of disulfide bond knots (Supplementary Figs. 2 and 3). The corresponding gene blocks were cloned into the pET24a vector using NdeI/NotI restriction sites. The assembled plasmids were transformed into *E. coli* BL21(DE3) cells (Thermo Fisher, EC0114) through heat shock transformation for 40 s at 42°C. For TMHC2, cells were cultured in 1 L of Luria Broth (LB) medium with 25 µg/mL kanamycin at 37°C until an OD_600_ of 0.8. For the ELP linker constructs, cells were cultured in 1 L of Terrific Broth (TB) medium with 25 µg/mL kanamycin at 37°C until an OD_600_ of 1.0. Protein overexpression was induced by adding 200 µM IPTG (for TMHC2) or 400 µM IPTG (for the ELP linker constructs), and incubated for 3 hr at 37°C. The cells were pelleted by centrifugation (5993×*g*, 10 min, 4°C) and resuspended in a lysis buffer: for TMHC2, 25 mM Tris-HCl, pH 7.4, 150 mM NaCl, 10% glycerol, 1 mM PMSF, 0.8 µg/ml pepstatin A, 0.8 µg/ml leupeptin, and 4 µg/ml aprotinin; for the ELP linker constructs, 50 mM Tris-HCl, pH 7.4, 200 mM NaCl, 1 mM TCEP, 10% glycerol, and 1 mM PMSF. Cell lysis was conducted using Emulsiflex-C3 at ∼17,000 psi. For TMHC2, additional step was added to extract membrane proteins—1% DDM detergent (GoldBio, DDM50) was added to the lysate and the sample was incubated for 1 hr at 4°C. Cell debris was removed by centrifugation (34,811×*g*, 30 min, 4°C), and the supernatant was incubated with 1 ml Ni-IDA resin (Clontech, 635662) for 1 hr at 4°C. The resin was packed into a column and washed with a wash buffer: for TMHC2, 25 mM Tris-HCl, pH 7.4, 150 mM NaCl, 10% glycerol, 0.1% DDM, and 20 mM imidazole; for the ELP linker constructs, 50 mM Tris-HCl, pH 7.4, 200 mM NaCl, 10% glycerol, and 20 mM imidazole. The protein was eluted with an elution buffer: for TMHC2, 25 mM Tris-HCl, pH 7.4, 150 mM NaCl, 10% glycerol, 0.1% DDM, and 300 mM imidazole; for the ELP linker constructs, 50 mM Tris-HCl, pH 7.4, 200 mM NaCl, 10% glycerol, and 300 mM imidazole. For TMHC2, further purification was performed using gel filtration (Cytiva, Superdex 200 10/30) in 25 mM Tris-HCl, pH 7.4, 150 mM NaCl, and 0.1% DDM. The purified samples were concentrated to ∼10 µM (for TMHC2) or ∼100 µM (for the ELP linker constructs) using Amicon Ultra-4 Centrifugal Filter Unit (3 kDa or 10 kDa, respectively; Merck), and then stored in aliquots at –80°C (Supplementary Fig. 2). For the Cys mutant of TMHC2, the same procedure as for TMHC2 was followed, except that the growth medium was switched to TB. Other proteins used in this work, such as mSpyCatcher and traptavidin, were also purified following formerly developed procedures^63,71^ (Supplementary Fig. 1).

### Molecular construct preparation for single-molecule tweezer experiments

2 µl of each ∼100 µM ELP linker construct was mixed with 10 µl of ∼10 µM TMHC2 and incubated for 3 hr at 25°C to form the TMHC2-ELP complex through SnoopTag-SnoopCatcher binding, with a covalent isopeptide bond forming spontaneously^79^. Two types of 1022-bp DNA handles were then attached to the TMHC2-ELP complex described below^63,70,71^. The DNA constructs, modified at one end with primary amine and at the other end with either azide or dual biotins (2×biotin), were generated using polymerase chain reaction (PCR) with λ DNA template (NEB, N3011S) and synthesized primers (IDT). The primers used in the PCR were a forward primer (ACAGAAAGCCGCAGAGCA) modified with amine at the 5′ end and a reverse primer (TCGCCACCATCATTTCCA) modified with azide or 2×biotin at the 5′ end. ∼8 ml of the PCR product (azide-DNA:2×biotin-DNA = 1:1) was purified using HiSpeed Plasmid Maxi Kit (Qiagen, 12663) and eluted with 1 ml of 0.1 M sodium bicarbonate, pH 8.3. For maleimide modification at the amine end of DNA using amine-NHS ester reaction, ∼1 µM of the eluted DNA sample was mixed with 4 µl of 250 mM SM(PEG)₂ (Thermo Scientific Pierce, 22102) and incubated for 20 min at ∼23°C. The sample was purified using Econo-Pac 10DG Desalting Column (Bio-Rad) and eluted with 1.5 ml of 0.1 M sodium phosphate, pH 7.3, 150 mM NaCl. To conjugate mSpyCatcher (with a cysteine) to the maleimide-modified DNA, ∼0.5 μM of the eluted DNA sample was mixed with 200 µl of ∼100 μM mSpyCatcher and incubated for 2 hr at ∼23°C. Free proteins were removed using HiTrap Q HP column (Cytiva, 17115401) with gradient elution from 0 to 1 M NaCl in 20 mM Tris-HCl (pH 7.5). The DNA peak fractions (∼30% of which corresponded to mSpyCatcher-conjugated DNA) were concentrated to ∼100 nM for the conjugated construct and stored at –80°C in 10 μl aliquots. The mSpyCatcher-conjugated DNA handles were then attached to the N-terminal ends of the monomers in the TMHC2-ELP complex through SpyTag-SpyCatcher binding, with a covalent isopeptide bond forming spontaneously^78^. To this end, 3 µl of the TMHC2-ELP complex sample was mixed with 10 µl of the mSpyCatcher-DNA sample and incubated for 2 hr at 25°C, yielding the final target construct (DNA-TMHC2-ELP complex; Supplementary Figs. 2 and 3). The mixture was diluted to ∼200 pM for the final target construct, with azide at one end and 2×biotin at the other end, and stored at –80°C in 10 μl aliquots. The dilution buffer was 25 mM Tris-HCl, pH 7.4, 150 mM NaCl, and 1.5% bicelle. The bicelles were composed of DMPC lipids (Avanti, 850345P) and CHAPSO detergents (Merck, C3649) at a 2.5:1 molar ratio^63,71^.

### Sample chamber preparation for single-molecule tweezer experiments

The coverslips (VWR, No. 1.5, 24×50 mm and 24×40 mm) were cleaned using KOH and Piranha solution^63,71^. The bottom coverslip (24×50 mm) was passivated with a polyethylene glycol (PEG) mixture of methyl-PEG (Laysan Bio, MPEG-SVA-5000) and biotin-PEG (Laysan Bio, Biotin-PEG-SVA-5000) at a 100:1 molar ratio^63,71^. These two surface-treated coverslips were assembled to construct a single-molecule sample chamber with a channel volume of ∼10 µl (1 CV). 1 µl of streptavidin-coated polystyrene beads (Spherotech, SVP-10-5) was washed and resuspended with 50 µl of 0.1 M sodium phosphate, pH 7.4, 150 mM NaCl, and 0.1% Tween 20. 1 CV of the polystyrene bead slurry was injected into the sample chamber and incubated for 2–5 min at 20– 22°C to allow for surface binding. These surface-bound polystyrene beads were used to correct for thermal drift of the sample chamber. 5 CV of 100 mg/ml bovine serum albumin (BSA) was injected into the chamber and incubated for 5 min at 20–22°C for further surface passivation. The sample chamber was washed with 5 CV of buffer A (50 mM Tris-HCl, pH 7.4, 150 mM NaCl, and 0.1% DDM). 10 µl sample containing ∼200 pM of the DNA-TMHC2-ELP complex was mixed with 1 µl of 0.04 µM traptavidin and incubated for 15 min at 20–22°C. 1 CV of this sample was injected into the chamber and incubated for 10 min at 20–22°C. To block unoccupied biotin-binding sites, 1 CV of a 30-nt biotin-labeled oligonucleotide (10 µM in buffer A) was injected into the chamber and incubated for 5 min at 20–22°C. The sample chamber was washed with 5 CV of buffer B (50 mM Tris-HCl, pH 7.4, 150 mM NaCl, and 0.05% DDM). Amine-coated magnetic beads (Thermo Fisher, 14307D) were modified with dibenzocyclooctyne (DBCO; Merck, 762040) using amine-NHS ester crosslinking chemistry^63,71^. 1 µl of the DBCO-coated magnetic beads was washed and resuspended in 20 µl buffer B. 1 CV of the magnetic bead slurry was injected into the chamber and incubated for 1 hr at 25°C. The magnetic beads were covalently attached to surface-tethered target constructs (DNA-TMHC2-ELP complex) via DBCO-azide conjugation^63,71^. The sample chamber was washed with 5 CV of buffer C (50 mM Tris-HCl, pH 7.4, 150 mM NaCl, and 1.3% bicelle). The bicelles were composed of DMPC lipids (Avanti, 850345P) and CHAPSO detergents (Merck, C3649) at a 2.5:1 molar ratio^63,71^.

### Magnetic tweezer instrumentation

Magnetic tweezers were constructed on inverted microscopes (Olympus, IX73) equipped with motorized XY stages (ASI, MS-2000FT or Prior Scientific, H117P1)^63,70,81^. Light-emitting diodes (Thorlabs, M455L4, λ = 447 nm) were used to illuminate the magnetic beads tethered to target molecular constructs. Three-dimensional positions of the tethered beads were tracked using a charge-coupled device camera (JAI, CM-040GE or CM-030GE) at a rate of 60–90 Hz. Changes in the extension (end-to-end distance) of the target molecule correspond to variations in bead height relative to the sample chamber surface, as indicated by changes in the diffraction pattern captured by the camera. Bead heights were calibrated by recording height-dependent diffraction patterns using a piezo nanopositioner (Mad City Labs, Nano-F100S), which moves the objective lens (Olympus, UPLFLN100XO2 or UPLXAPO100XO) in precise increments to adjust the bead’s diffraction patterns. Real-time tracking of bead height was achieved using chi-square analysis based on this calibration data. The non-magnetic polystyrene beads immobilized on the chamber surface were also tracked to correct for thermal drift of the sample chamber. Mechanical force was applied to the target molecule using a pair of 10×10×12 mm neodymium magnets in an antiparallel configuration with a 1-mm gap. Vertical and rotational movements of the magnets were controlled using a translational motor (PI, M-126.PD1 or M-126.PD2) and a rotational motor (PI, DT-50 or DT-34). The applied force was calibrated as a function of the magnet position using an inverted pendulum model^63,81^. The magnetic tweezer setups were mounted on pneumatic optical tables (Daeil Systems, DVIO-I-1512M-300t800H) in imaging rooms maintained at 20–22°C.

### Single-molecule tweezer experiments

In the force-cycle experiments, mechanical force was gradually applied and released on the target molecular construct, with the following parameters: a force range of 1–50 pN, a magnet speed (*m*_s_) of 0.1 mm/s, and a 1-s pause at either 1 or 50 pN. In our magnetic tweezer setup, the force-loading rate ranges from 0.035 to 3.1 pN/s across the force span of 1–50 pN at *m*_s_ = 0.1 mm/s. The force-extension curves were median-filtered for extension and smoothed for force, each using a window size of 10. The distributions of dissociation forces and step sizes in each ELP linker system were analyzed using the worm-like chain (WLC) model, represented by the Marko–Siggia equation^72,82^, *FL*_p_/*k*_B_*T* = *l*/*L*_c_ + (1–*l*/*L*_c_)^−2^/4 – 1/4. In this equation, *F* is the applied force, *k*_B_ is the Boltzmann constant, *T* is the absolute temperature, *l* is the extension of the ELP linkers, *L*_p_ is the persistence length of the ELP linkers (measured as 0.35 nm; ref. ^59^), and *L*_c_ is the contour length of the ELP linkers, derived by multiplying the number of bonds between residues by the average bond length of 0.36 nm (refs. ^63,83^). The probability of dimerization in the force-cycle experiments in Fig. 2e was estimated as the proportion of force-extension traces showing dissociation events for each molecule. These force-cycle experiments were performed in the three EPL systems with varying linker lengths. In the force-clamp experiments, a specified force level is driven by a rapid magnet speed of *m*_s_ = 15 mm/s and then maintained as constant during a specified time span. Two types of force clamp experiments were conducted: a high-force or low-force clamp mode focused on characterizing the intermediate states and their transition kinetics during the dissociation or dimerization process, respectively (see the following sections for details on relevant analyses). In the high-force clamp mode, a constant 19–20 pN was applied to the molecular construct and held for up to 3 min until monomer dissociation occurred. Following dissociation, the force was reduced back to 1 pN and held for 60 s to allow re-dimerization of monomers before beginning the next cycle. These high-force clamp experiments were performed on both the wild-type and Cys mutant of TMHC2 in the EPL-L linker system. In the low-force clamp experiments, the TMHC2 dimer was first fully dissociated into its monomers at 35 pN for 20 s (Supplementary Fig. 9c). Following dissociation, the force was reduced to a constant 5–13 pN and held for 3 min to allow re-dimerization of monomers before beginning the next cycle. These low-force clamp experiments were performed in the three EPL systems with varying linker lengths. The probability of dimerization up to each state in the force-clamp experiments was estimated as the proportion of time-resolved extension traces showing dimerization up to each state (Figs. 3b and 6g). To examine the effects of peptide competition, two types of peptides, TRTEIIR (peptide I) and TRTEIIRELERSL (peptide II), were synthesized by Peptron (South Korea) (Supplementary Fig. 4c). For the single-molecule tweezer experiments, 500 µM peptide stocks in deionized water (DW) were diluted to 10 µM using 1.3% bicelle (see the previous sections for details on bicelle composition). The same force-cycle and -clamp experiments and the relevant analyses were conducted following the injection of 5 CV peptide solution into the sample chamber. The probability of complete dimerization in the force-cycle experiments in Fig. 6c was estimated as the proportion of force-extension traces showing normal dissociation forces and step sizes consistent with the WLC model (Supplementary Fig. 25).

### Analysis of dimerization intermediate states

The force-clamp extension traces were median-filtered using a 5-Hz window. The distribution of extension values was modeled using the Gaussian mixture model (GMM). The Bayesian information criterion (BIC) was used to determine the optimal number of states in the traces, defined as BIC = *q*ln(*n*) − 2ln(*L̂*), where *q* is the number of output parameters given by the GMM, *n* is the number of data points, and *L̂* is the maximized value of the likelihood function of the GMM^83–85^. The optimal number of states was determined from the BIC plot as the elbow point, where the slope of the BIC changes substantially with the number of states^83,85^ (Supplementary Fig. 14). The extension position for each state was determined using the hidden Markov model (HMM), with the initial positions derived from the GMM^63,84^. The Baum-Welch algorithm, a specific form of the expectation–maximization (EM) algorithm, was used to iteratively maximize the likelihood and optimize the HMM model parameters^63,84^. The most probable state at each time point was assigned using the Viterbi algorithm, and the dwell times in one state before transitioning to another state were calculated. Using the identified extension position for each state, the step sizes for intermediate states (*I*_1_ and *I*_2_) during dimerization or dissociation were normalized to the total step sizes between the monomer (*M*) and dimer (*D*) states, with values of 0 and 1 assigned to the *M* and *D* states, respectively. The normalized step sizes for the *I*_1_ and *I*_2_ states were determined as 0.19±0.05 (SD) and 0.78±0.08 (SD), respectively, and were mapped onto the protein structure based on the designated 0^th^–10^th^ interaction layers. The 1^st^–10^th^ layers correspond to the leucine zipper-like, closely interacting inner residues, appearing every 3–4 residues, while the 0^th^ layer corresponds to the flexible polypeptide linker residues connecting monomer helices, allowing close interactions between the monomers (Supplementary Fig. 4a,b).

### Characterization of transition kinetics between states

The transition rate constants as a function of force (*k*_a_ for association, *k*_d_ for dissociation) were obtained from the mean dwell times in one state before transitioning to another state at constant 5–13 pN (Supplementary Fig. 22). The *k*_a_ and *k*_d_ values were extrapolated to the rate constants at zero force (*k*_a0_, *k*_d0_) using the Bell-Evans model^72,86–88^: *k* = *k*_0_·exp(±*F*Δ*x*^†^/*k*_B_*T*) (+ for *k*_d_ and – for *k*_a_), where *F* is the applied force, Δ*x*^†^ is the distance to the transition state, *k*_B_ is the Boltzmann constant, and *T* is the absolute temperature. The post-diffusion kinetics at zero force were then obtained by correcting for the ELP linker length. To this end, the *k*_a0_ and *k*_d0_ values, as a function of the ELP linker extension at zero force (*d*_elp_), were extrapolated to the rate constants at the minimum distance between monomers (*d*_min_), denoted as *k*_a0,m_ and *k*_d0,m_. The *d*_elp_ and *d*_min_ values were estimated from the root mean square end-to-end distance of the WLC polymer, *d*_rmsd_ = 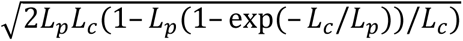 (refs. ^89–91^), and the dimer’s structural information (Supplementary Fig. 23). In the *d*_rmsd_ formula, *L*_p_ is the persistence length of the ELP linkers (measured as 0.35 nm; ref. ^59^), and *L*_c_ is the contour length of the ELP linkers, derived by multiplying the number of bonds between residues by the average bond length of 0.36 nm (refs. ^63,83^). The estimated *d*_elp_ and *d*_min_ values were *d*_elp-l_ = 6.02 nm, *d*_elp-m_ = 5.24 nm, *d*_elp-s_ = 4.29 nm, and *d*_min_ = 3.84 nm (Supplementary Fig. 23). The Δ*x*^†^ values as a function of *d*_elp_ were also extrapolated to those at *d*_min_, denoted as Δ*x*^†^_m_, which were used to reconstruct the free energy landscape for the post-diffusion dimerization (see the following section). The rate constants for complete dissociation from the *D* to *M* state were also determined from both the force-clamp and force-cycle experiments. In the force-clamp experiments, the rate constants for complete dissociation (*k*_d,com_) as a function of force were obtained from the distributions of the total durations before transitioning from the *D* to *M* state at constant 20–30 pN (Supplementary Fig. 9b–d). To obtain the rate constant for complete dissociation at zero force (*k*_d0,com_), the *k*_d,com_ values were extrapolated to zero force using the Bell-Evans model, as described above (Supplementary Fig. 9d,e). In the force-cycle experiments, the cumulative plots of dissociation force distributions were normalized to obtain the dissociation probabilities as a function of force (Supplementary Fig. 9a). To obtain *k*_d0,com_, the dissociation probability profiles were fitted with an equation derived from the first-order rate equation and the Bell-Evans equation: *P*_d,com_ = 1–exp(∫d*F*(– *k*_d0,com_·exp(*F*Δ*x*^†^_d,com_/*k*_B_*T*)/*Ḟ*)) (ref. ^63^), where *P*_d,com_ is the probability for complete dissociation from the *D* to *M* state, *F* is the force, *Ḟ* is the force-loading rate, *k*_d0,com_ is the rate constant for complete dissociation at zero force, and Δ*x*^†^_d,com_ is the distance to the transition state.

### Reconstruction of dimerization energy landscapes

The free energy landscape for the post-diffusion dimerization process was reconstructed using the kinetic estimates corrected for the ELP linker length (*k*_0,m_ and Δ*x*^†^_m_) and the characterized intermediate positions (Supplementary Fig. 24). The energy barrier heights between the states (Δ*G*^†^_a,m_ and Δ*G*^†^_d,m_) were obtained using the Kramers rate framework^75^: *k*_0,m_ = *k*_ω_·exp(–Δ*G*^†^_m_/*k*_B_*T*), where *k*_ω_ is the frequency factor, *k*_B_ is the Boltzmann constant, and *T* is the absolute temperature. Different estimates of *k*_ω_ were used for different dimerization domains, reflecting its dependence on solvent viscosity^63,76^—*k*_ω_ = 10^4^–10^6^ s^−1^ for the soluble domain exposed to water^77^ and *k*_ω_ = 45.15±3.56 (SE) s^−1^ for the TM domains embedded in lipid bilayers^63^. The reaction coordinate was defined as the extent of dimerization from the *M* to *D* state across the designated 0^th^–10^th^ interaction layers, with values of 0 and 1 assigned to the *M* and *D* states, respectively. The positions of the intermediate states and transition states were mapped onto the reaction coordinate based on the normalized dimerization step sizes and the normalized Δ*x*^†^_a,m_ and Δ*x*^†^_d,m_ values, respectively. For comparison, the dimerization energy landscape in each ELP linker system (without linker correction) was also derived from the kinetic estimates (*k*_0_ and Δ*x*^†^) and the intermediate positions in each system (Supplementary Fig. 24).

### Transmission electron microscopy (TEM)

Negative staining TEM was utilized to visualize bicelles and measure their sizes. A solution of 1.3% (w/v) DMPC bicelles, used in the single-molecule tweezer experiments, was applied onto a formvar/carbon-coated copper grid (Ted Pella, 01754-F). After 1 min, excess sample was blotted off using a filter paper. The grid was gently rinsed with distilled water and stained with 2% ammonium molybdate tetrahydrate for 10 s. Excess staining solution was removed using a filter paper, and the grid was left to air dry for 5 min. TEM images were obtained using JEM-1400 TEM (JEOL, Japan) operating at 120 kV. The images were analyzed using ImageJ software to determine the diameter of the bicelles.

### All-atom molecular dynamics (AAMD) simulation

The AAMD simulation system was constructed using CHARMM-GUI Membrane Builder^92^. The protein structure of the TMHC2 dimer was obtained from AlphaFold3^62,93^, and the dimer was inserted into the lipid bilayer composed of 302 DMPC molecules, using coordinates optimized by the PPM 2.0 (ref. ^94^). The simulation system was solvated with TIP3P water molecules^95^, adjusted to pH 7.0, and neutralized with ∼150 mM K^+^ and Cl^−^ ions. The final dimensions of the system were 10.1×10.1×10.5 nm, containing a total of 100,850 atoms. Periodic boundary conditions were applied to the simulation system. The MD simulations were conducted using GROMACS (version 2021.4) with CHARMM36m force field (charmm36-mar2019)^96,97^. The simulation system was energy-minimized using the steepest descent algorithm. During the energy minimization, van der Waals (vdW) interactions were calculated with a cutoff distance of 12 Å, smoothly switching off at 10–12 Å using a force-switch function^98^. Long-range electrostatic interactions were calculated using the Particle-Mesh Ewald (PME) method with a cutoff distance of 12 Å (ref. ^99^). The LINCS algorithm was employed to impose constraints on the bond lengths involving hydrogen atoms^100^. Following the energy minimization, the system underwent a multi-stage equilibration process. Initially, the system equilibration was performed in the NVT ensemble for a total of 375,000 steps with a 1-fs time step. The temperature was maintained at 303.15 K using the Berendsen thermostat^101^. The equilibration continued in the NPT ensemble for an additional 750,000 steps with a 2-fs time step. The pressure was maintained at 1 bar using the Berendsen barostat with semi-isotropic pressure coupling and a compressibility of 4.5 × 10⁻⁵ bar⁻¹ (ref. ^101^). The LINCS method was applied to handle the bond constraints throughout both the NVT and NPT equilibration steps. After the equilibration, a production run was performed for 1 μs with a 2-fs time step. During the production run, the temperature was maintained at 303.15 K using the Nosé-Hoover thermostat^102^, and the pressure was maintained at 1 bar using the Parrinello-Rahman barostat with isotropic pressure coupling and a compressibility of 4.5 × 10⁻⁵ bar⁻¹ (ref. ^103^). The 1-μs MD simulation of the TMHC2 dimer was analyzed using GROMACS modules, the Residue Interaction Network Generator (RING)^73,74^, and Python.

### Analysis of AAMD simulation trajectories

The analysis of solvent environments around the TMHC2 dimer, as shown in Fig. 3h, was conducted using a 1-ns time interval trajectory of a 1-μs AAMD simulation of the dimer. The center of the dimer was translated to the center of the simulation system using the GROMACS trjconv module. The simulation trajectory was analyzed using MDAnalysis library in Python^104^. The z-coordinates of atoms within a 2-nm radius cylinder centered on the dimer’s center of mass were extracted at each time frame. The analyzed atoms include those of water molecules, lipid heads with glycerol backbone, lipid tails, and the closely interacting protein residues at the dimerization interface. The layers of closely interacting protein residues (the interaction layers) are shown in Supplementary Fig. 4a,b. The z-coordinates of the atoms were binned into 1 Å intervals, and the z-coordinate counts for each type of molecules and each interaction layer of the protein were normalized by their maximum counts. The distributions of z-coordinate counts for solvent components (water, lipid heads, and lipid tails) were compared with those for the protein’s interaction layers (Fig. 3h). The intermolecular interactions between monomers in the dimer state, as shown in Fig. 3g, were also analyzed using the AAMD simulation trajectory. At each time frame, the interaction types and counts for residue pairs between the monomers were obtained using the residue interaction network generator (RING) (version 4.0; https://ring.biocomputingup.it; performed in July 2024)^73,74^. The distance thresholds for each type of interaction used in the RING analysis were as follows: 3.9 Å for hydrogen bonds between donor and acceptor, 2.5 Å for hydrogen bond between hydrogen and acceptor, 4.3 Å for π–hydrogen bond between donor and an aromatic ring center, 6.5 Å for π–π interactions between aromatic ring centers, 5 Å for π–cation interactions between an aromatic ring and the cation center of mass, 4 Å for ionic bonds (salt bridges), 2.5 Å for disulfide bonds, 2.8 Å for metal ion coordination between metal ion atoms and acceptor, and 0.01 for vdW interactions as the fraction of intersection between the vdW radii of two atoms. In this analysis, three types of intermolecular interactions—hydrogen bonds, ionic interactions, and vdW interactions—were detected between the monomers. For each interaction type, the interaction counts were summed for each interaction layer region over 1 μs (Supplementary Fig. 19 for the detailed analysis procedure).

### Coarse-grained molecular dynamics (CGMD) simulation

Two CGMD simulation systems were prepared as follows: 1) the initial dimer system, where the TMHC2 dimer was embedded within the lipid bilayer, and 2) the initial monomer system, where the two monomers of TMHC2 were initially separated by ∼10 nm within the lipid bilayer. To this end, AAMD simulation systems with the protein/membrane configurations were constructed using CHARMM-GUI Membrane Builder^92^, and then converted into the coarse-grained (CG) representation using CHARMM-GUI Martini Maker with MARTINI 3.0 force field^105,106^. The lipid membranes consisted of 304 and 897 DPPC molecules for the initial dimer and monomer systems, respectively. The CGMD simulation systems were solvated with MARTINI polarized water molecules^107^, adjusted to pH 7.0, and neutralized with ∼150 mM Na^+^ and Cl^−^ ions. The final dimensions of the systems were 10.0×10.0×10.2 nm with 8,679 beads for the initial dimer system and 17.0×17.0×12.9 nm with 31,305 beads for the initial monomer system, respectively. Periodic boundary conditions were applied to both simulation systems. The CGMD simulations were conducted using GROMACS (version 2021.4) with MARTINI 3.0 force field^106^. The simulation systems were energy-minimized using the steepest descent algorithm, with vdW interactions calculated with a cutoff distance of 1.1 nm. Long-range electrostatic interactions were handled using the reaction-field method with a dielectric constant of 15 and a 1.1-nm cutoff^108^. Temperature coupling was controlled using the v-rescale thermostat with a time constant of 1.0 ps (ref. ^109^), maintaining the systems at 303.15 K. Semi-isotropic pressure coupling was applied using the Berendsen barostat with a time constant of 5.0 ps and a compressibility of 3 × 10⁻4 bar⁻¹ (ref. ^101^). Following the energy minimization, the systems underwent a multi-stage equilibration process. Initially, the system was equilibrated in the NPT ensemble for 500,000 steps with a 2.0-fs time step, under the same temperature, pressure, and interaction conditions as in the energy minimization stage. As equilibration progressed, the time step was incrementally increased from 2.0 fs to 20.0 fs, resulting in a total equilibration time of 14 ns under periodic boundary conditions. After the equilibration stage, production runs were performed in the NPT ensemble for 2 μs with a 20-fs time step. The temperature was maintained at 303.15 K using v-rescale thermostat^109^, while the pressure was controlled at 1 bar using the semi-isotropic Parrinello-Rahman barostat with a compressibility of 3 × 10⁻4 bar⁻¹ (ref. ^103^). The 2-μs CGMD simulations were analyzed using GROMACS modules and Python.

### Analysis of CGMD simulation trajectories

The dimerization percentage of TMHC2 as a function of time and the binding order across the interaction layers were analyzed from the CGMD simulation trajectories. The distance cutoff for binding at the interaction layers (*d*_cut,0_) was initially determined from five independent trajectories of the initial dimer system, as described below. For each trajectory, the backbone bead coordinates of each layer’s residues in two monomers were extracted at 1-ns intervals and averaged to obtain their center coordinates. The distance between the center coordinates in each layer (*d_mm_*) was then calculated. The proportion of all layer’s *d_mm_* values below an arbitrary distance cutoff *d*_cut_ (*P*_dimer_; dimerization percentage) was plotted as a function of *d*_cut_ (Supplementary Fig. 20). From the plot of *P*_dimer_ *vs d*_cut_, the *d*_cut_ at *P*_dimer_ = 99% was selected as *d*_cut,0_ (17.3 Å ; Supplementary Fig. 20). For a hundred independent trajectories of the initial monomer system, the same *d*_cut,0_ criterion was applied to obtain *P*_dimer_ as a function of time, as shown in Fig. 4d,e. Only 80 trajectories showed dimerization events within the 2-μs time span, and only these trajectories were considered for further analysis. The period of the post-diffusion dimerization was defined as the interval from the last time point at *P*_dimer_ = 0% to the first time point at *P*_dimer_ = 100%. The criterion for determining the dimerized states in the initial monomer system was the first time point at *P*_dimer_ = 100%. The time-resolved binding profiles at each interaction layer, as shown in Fig. 4f,g, were also determined based on the same distance cutoff, *d*_cut,0_. The binding order across the layers was further estimated using two different criteria: the first binding time and the total bound state durations during the post-diffusion dimerization period (Supplementary Fig. 21). With each criterion, all possible binding pathways across the layers were identified. The binding order across the layers was then quantified as the relative frequency of the rank order at each layer over all 80 trajectories that exhibited dimerization events.

## Acknowledgments

This work was supported by the National Research Foundation of Korea (2020R1C1C1003937 to D.M. and 2021R1C1C1010943 to J.M.C.).

## Author Contributions

D.M. conceived and supervised the project. V.W.S., S.K., and D.M. designed the single-molecule tweezer experiments. V.W.S. performed the single-molecule tweezer experiments. V.W.S., S.K., E.K., and D.M. analyzed the single-molecule experiment data. V.W.S. and S.K. purified the proteins and prepared the molecular constructs. E.K. and D.M. performed the MD simulations. E.K., T.S.L., J.M.C., and D.M. analyzed the MD simulation data. W.C.B.W. performed the TEM measurements. V.W.S., E.K., S.K., and D.M. prepared the manuscript with input from all authors.

## Data Availability

All data that support the findings of this study are available in the main text, Supplementary Information, and Source Data, as well as from the corresponding author upon request. Other raw data such as the MD simulation trajectory files have been deposited to figshare, which are available in [https://doi.org/10.6084/m9.figshare.28218593]. Source data are provided with this paper.

## Code Availability

Analysis codes are available in Code Ocean [https://doi.org/10.24433/CO.9572266.v1].

