## Supplementary Information for "Single-molecule tweezers decoding hidden dimerization patterns of membrane proteins within lipid bilayers"

### Contents


Supplementary Fig. 1 | Amino acid sequences of proteins
Supplementary Fig. 2 | Electrophoresis for molecular constructs and conjugations
Supplementary Fig. 3 | Molecular construct and force propagation
Supplementary Fig. 4 | Protein interaction layers and peptides
Supplementary Fig. 5 | Dissociation forces and step sizes of TMHC2 dimer
Supplementary Fig. 6 | Bicelle size characterized using TEM
Supplementary Fig. 7 | TMHC2 dimerization within lipid bilayers
Supplementary Fig. 8 | Representative high force traces in ELP-L system
Supplementary Fig. 9 | Transition kinetics for complete dissociation of TMHC2 dimer
Supplementary Fig. 10 | Representative low force traces in ELP-S system
Supplementary Fig. 11 | Representative low force traces in ELP-M system
Supplementary Fig. 12 | Representative low force traces in ELP-L system
Supplementary Fig. 13 | Representative low force traces of Cys mutant in ELP-L system
Supplementary Fig. 14 | Bayesian information criterion (BIC) analysis
Supplementary Fig. 15 | Transition step sizes of TMHC2
Supplementary Fig. 16 | Transition step sizes of Cys mutant or WT with peptide II
Supplementary Fig. 17 | Probability of dimerization in force-clamp experiments
Supplementary Fig. 18 | Normalized step sizes up to each intermediate state
Supplementary Fig. 19 | Analysis procedure of intermolecular interactions
Supplementary Fig. 20 | Determination of distance cutoff for binding at interaction layers
Supplementary Fig. 21 | Analysis of binding order across interaction layers
Supplementary Fig. 22 | Force-dependent transition kinetics
Supplementary Fig. 23 | ELP linker extension at zero force and minimum monomer distance
Supplementary Fig. 24 | Reconstructed energy landscapes of TMHC2 dimerization
Supplementary Fig. 25 | Dissociation forces and step sizes in the presence of peptides
Supplementary Fig. 26 | Representative low force traces in the presence of peptide II

Supplementary Table 1 | Number of transitions in force-clamp experiments
Supplementary Table 2 | Kinetic and energetic estimates at zero force

**TMHC2 monomer with SpyTag and SnoopTag**

SpyTag SnoopTag TMHC2 monomer

MHHHHHHGGSAHIVMVDAYKPTKGGSGGSKLGDIEFIKVNKGGSGGSMTRTEIIRELERSLRLQLVLAIFLMALLIVLLWLQQNGSSNNNVNYLLIVILVL
VLVIVALAVTQKYLVEQLKRQDLE

R3C/D76C for Cys mutant

**ELP linkers with SnoopCatchers**

— Poypeptide region including ELP linker
C: Cysteines that form an internal disulfide bond

SnoopCatcher ELP-S

MHHHHHHGGSKPLRGAVFSLQKQHPDYPDIYGAIDQNGTYQNVRTGEDGKLTFKNLSDGKYRLFENSEPAGYKPVQNKPIVAFQIVNGEVRDVTSIVP
QDIPATYEFTNGKHYITNEPIPPKGGSGGSDGSGGSGGQLDGHGVGVPGVGVPGVGVPGEGVPGVGVPGVGVPGVGVPGVGVPGEGGGSGGSCV
PKPLRGAVFSLQKQHPDYPDIYGAIDQNGTYQNVRTGEDGKLTFKNLSDGKYRLFENSEPAGYKPVQNKPIVAFQIVNGEVRDVTSIVPQDIPATYEFTN
GKHYITNEPIPPKC

SnoopCatcher

ELP-M

MHHHHHHGGSKPLRGAVFSLQKQHPDYPDIYGAIDQNGTYQNVRTGEDGKLTFKNLSDGKYRLFENSEPAGYKPVQNKPIVAFQIVNGEVRDVTSIVP
QDIPATYEFTNGKHYITNEPIPPKGGSGGSDGSGGSGGQLDGHGVGVPGVGVPGVGVPGEGVPGVGVPGVGVPGVGVPGVGVPGEGVPGVGVPGV
GVPGVGVPGVGVPGEGVPGVGVPGVGEGGSGGSCVPKPLRGAVFSLQKQHPDYPDIYGAIDQNGTYQNVRTGEDGKLTFKNLSDGKYRLFENSEPA
GYKPVQNKPIVAFQIVNGEVRDVTSIVPQDIPATYEFTNGKHYITNEPIPPKC

ELP-L

MHHHHHHGGSKPLRGAVFSLQKQHPDYPDIYGAIDQNGTYQNVRTGEDGKLTFKNLSDGKYRLFENSEPAGYKPVQNKPIVAFQIVNGEVRDVTSIVP
QDIPATYEFTNGKHYITNEPIPPKGGSGGSDGSGGSGGQLDGHGVGVPGVGVPGVGVPGEGVPGVGVPGVGVPGVGVPGVGVPGEGVPGVGVPGV
GVPGVGVPGVGVPGEGVPGVGVPGVGELYAVTGRGRAPASSAPIATSVPGVGVPGVGVPGEGGGSGGSCVPKPLRGAVFSLQKQHPDYPDIYGAID
QNGTYQNVRTGEDGKLTFKNLSDGKYRLFENSEPAGYKPVQNKPIVAFQIVNGEVRDVTSIVPQDIPATYEFTNGKHYITNEPIPPKC

**mSpyCatcher**

MBP

C: Cysteine for DNA attachment

MKIEEGKLVIWINGDKGYNGLAEVGKKFEKDTGIKVTVEHPDKLEEKFPQVAATGDGPDIIFWAHDRFGGYAQSGLLAEITPDKAFQDKLYPFTWDAVR
YNGKLIAYPIAVEALSLIYNKDLLPNPPKTWEEIPALDKELKAKGKSALMFNLQEPYFTWPLIAADGGYAFKYENGKYDIKDVGVDNAGAKAGLTFLVDLI
KNKHMNADTDYSIAEAAFNKGETAMTINGPWAWSNIDTSKVNYGVTVLPTFKGQPSKPFVGVLSAGINAASPNKELAKEFLENYLLTDEGLEAVNKDK
PLGAVALKSYEEELAKDPRIAATMENAQKGEIMPNIPQMSAFWYAVRTAVINAASGRQTVDEALKDAQTNGMSYYHHHHHHDYDIPTTENLYFQGGC
GDTLSGLSSEQGQSGDMTIEEDSATHIKFSKRDEDGKELAGATMELRDSSGKTISTWISDGQVKDFYLYPGKYTFVETAAPDGYEVATAITFTVNEQG
QVTVNGKATKGDAHI

SpyCatcher

TEV protease cleavage site

**Traptavidin**

MAEAGITGTWYNQLGSTFIVTAGADGALTGTYESAVGNAEGDYVLTGRYDSAPATDGSGTALGWTVAWKNNYRNAHSATTWSGQYVGGAEARINTQ
WLLTSGTTEANAWKSTLVGHDTFTKVKPSAASGGSHHHHHH

**Supplementary Fig. 1 | Amino acid sequences of proteins.** SpyTag and SnoopTag on TMHC2 spontaneously bind to the SpyCatcher protein of mSpyCatcher and the SnoopCatcher protein of elastin-like polypeptide (ELP) linker constructs, respectively, forming isopeptide bonds. The cysteine positions in the TMHC2 Cys mutant are underlined. The cysteines flanking the C-terminal SnoopCatcher in the ELP linker constructs form disulfide bonds to prevent its unfolding (see Supplementary Fig. 3 for details). The polypeptide regions, including the ELP linkers, are 75, 111, and 146 aa for the ELP-S, -M, and -L linker constructs, respectively (underlined). GGS linkers are inserted between the proteins, peptide tags, and ELP linkers. 6xHis tags are used to purify the proteins. The cystine between MBP and SpyCatcher in mSpyCatcher is used to attach the protein to DNA handles. The sequence of the TEV protease cleavage site in mSpyCatcher is underlined. Traptavidin is used to anchor the final single-molecule constructs to the sample chamber surface.

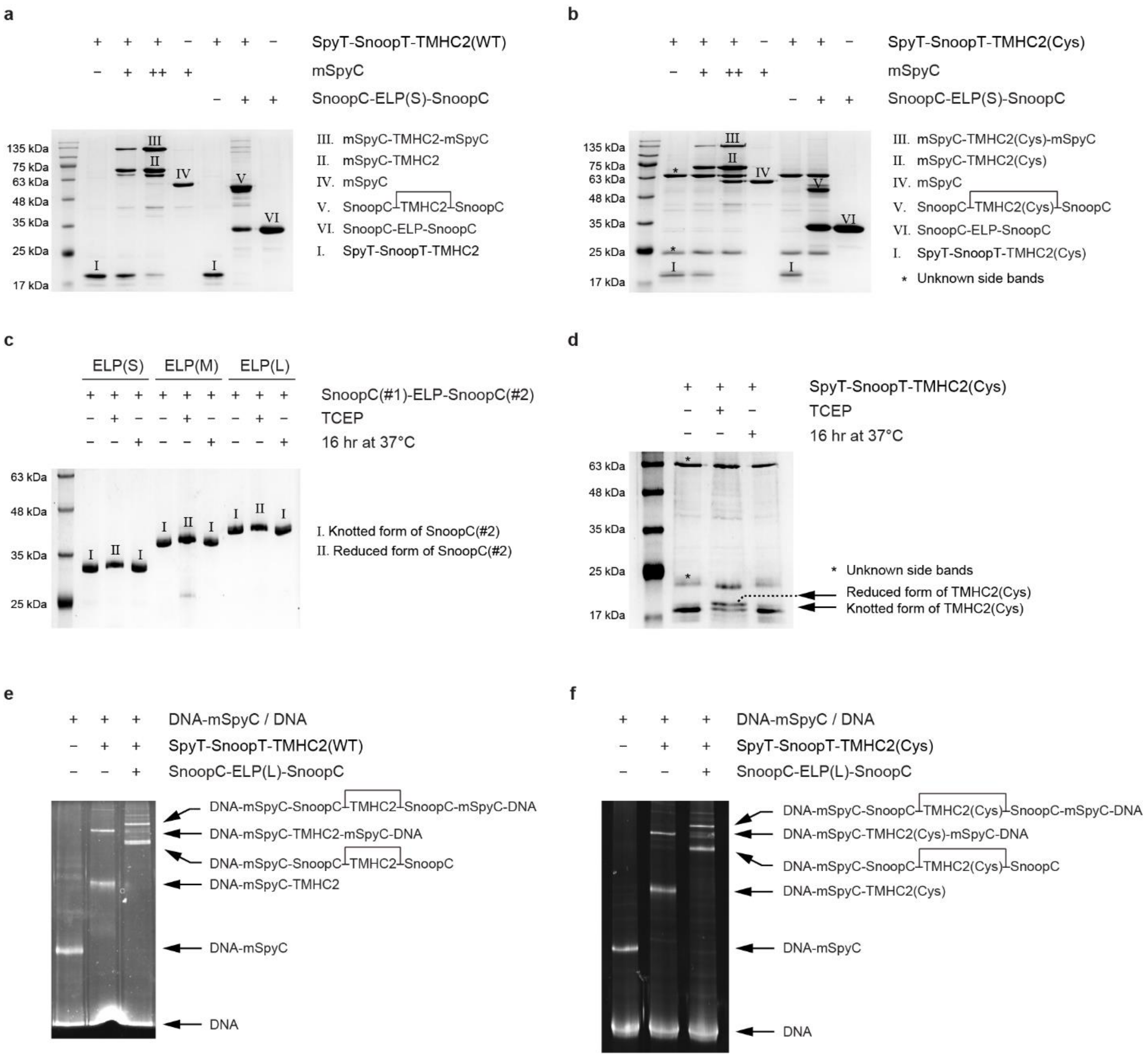


**Supplementary Fig. 2 | Electrophoresis for molecular constructs and conjugations.** (**a**,**b**) Protein purification and conjugation confirmed by 12% SDS–PAGE. The wild-type (WT) and Cys mutant of TMHC2 are shown in panels **a** and **b**, respectively. Successful protein conjugations mediated by SpyT-SpyC or SnoopT-SnoopC binding are tested upon protein purification. The notation "++" indicates a higher concentration of mSpyC. The leftmost lanes represent protein markers. (**c**,**d**) Disulfide formation within proteins confirmed by 12% SDS–PAGE (non-reducing). The disulfide bonds are formed within SnoopC(#2) of the ELP constructs (**c**) and within the monomers of the TMHC2 Cys mutant (**d**). Refer to Fig. 2h (inset) and Supplementary Figs. 1 and 3 for more details. TCEP is a reducing agent that converts disulfide bonds into cysteines. Prolonged incubation at a higher temperature of 37°C facilitates disulfide bond formation. The leftmost lanes represent protein markers. (**e**,**f**) Conjugation of TMHC2, ELP construct, and DNA handle confirmed by 6% SDS–PAGE. The wild-type (WT) and Cys mutant of TMHC2 are shown in panels **e** and **f**, respectively. The conjugation of TMHC2 with the DNA handle is mediated by SpyT-SpyC binding, while its conjugation with the ELP construct is mediated by SnoopT-SnoopC binding. The conjugated constructs, with azide at one end and 2×biotin at the other end, can only be tethered to both the sample chamber surface and the magnetic beads for single-molecule tweezer experiments. Abbreviations: ELP, elastin-like polypeptide linker; ELP(S), ELP with a short length; ELP(M), ELP with a middle length; ELP(L), ELP with a long length; SpyT, SpyTag; SnoopT, SnoopTag; SpyC, SpyCatcher; mSpyC, MBP-fusion SpyCatcher; SnoopC, SnoopCatcher; SnoopC(#1), N-terminal SnoopC in SnoopC-ELP-SnoopC constructs; SnoopC(#2), C-terminal SnoopC in SnoopC-ELP-SnoopC constructs; SDS–PAGE, sodium dodecyl sulfate–polyacrylamide gel electrophoresis; TCEP, tris(2-carboxyethyl)phosphine.

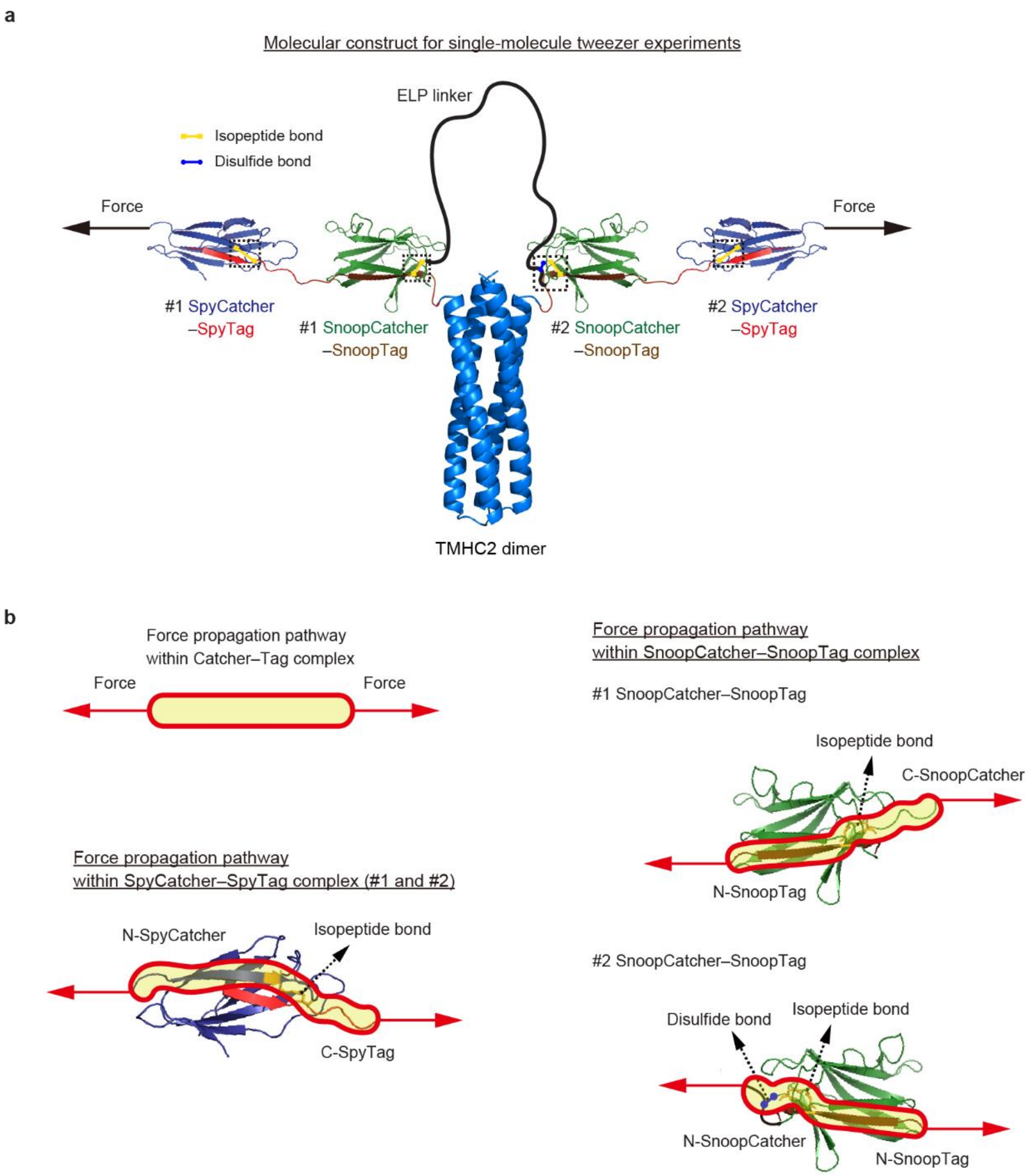


**Supplementary Fig. 3 | Molecular construct and force propagation.** (**a**) Molecular construct for single-molecule tweezer experiments. SpyTag and SnoopTag are genetically linked to the N-terminal end of TMHC2 monomers. The ELP linker flanked by SnoopCatcher proteins is attached to the TMHC2 dimer through SnoopCatcher-SnoopTag binding. The DNA handle-conjugated SpyCatcher is attached to each TMHC2 monomer through SpyCatcher-SpyTag binding. Once bound, internal isopeptide bonds within the Catcher-Tag complexes spontaneously form and represented by yellow lines. The disulfide bond in the #2 SnoopCatcher-SnoopTag complex, involving the C-terminal SnoopCatcher of the ELP linker construct, is represented by a blue line.

The isopeptide and disulfide bonds are highlighted with dashed boxes. (**b**) Force propagation pathways within the Catcher-Tag complexes. In each Catcher-Tag complex, the force is expected to propagate along the shortest pathway involving the denoted isopeptide and disulfide bonds, thereby preventing the global unfolding of the complexes. The N- or C-terminus of the Catcher proteins and peptide Tags is indicated by abbreviations, such as N-SpyCatcher and C-SpyTag.

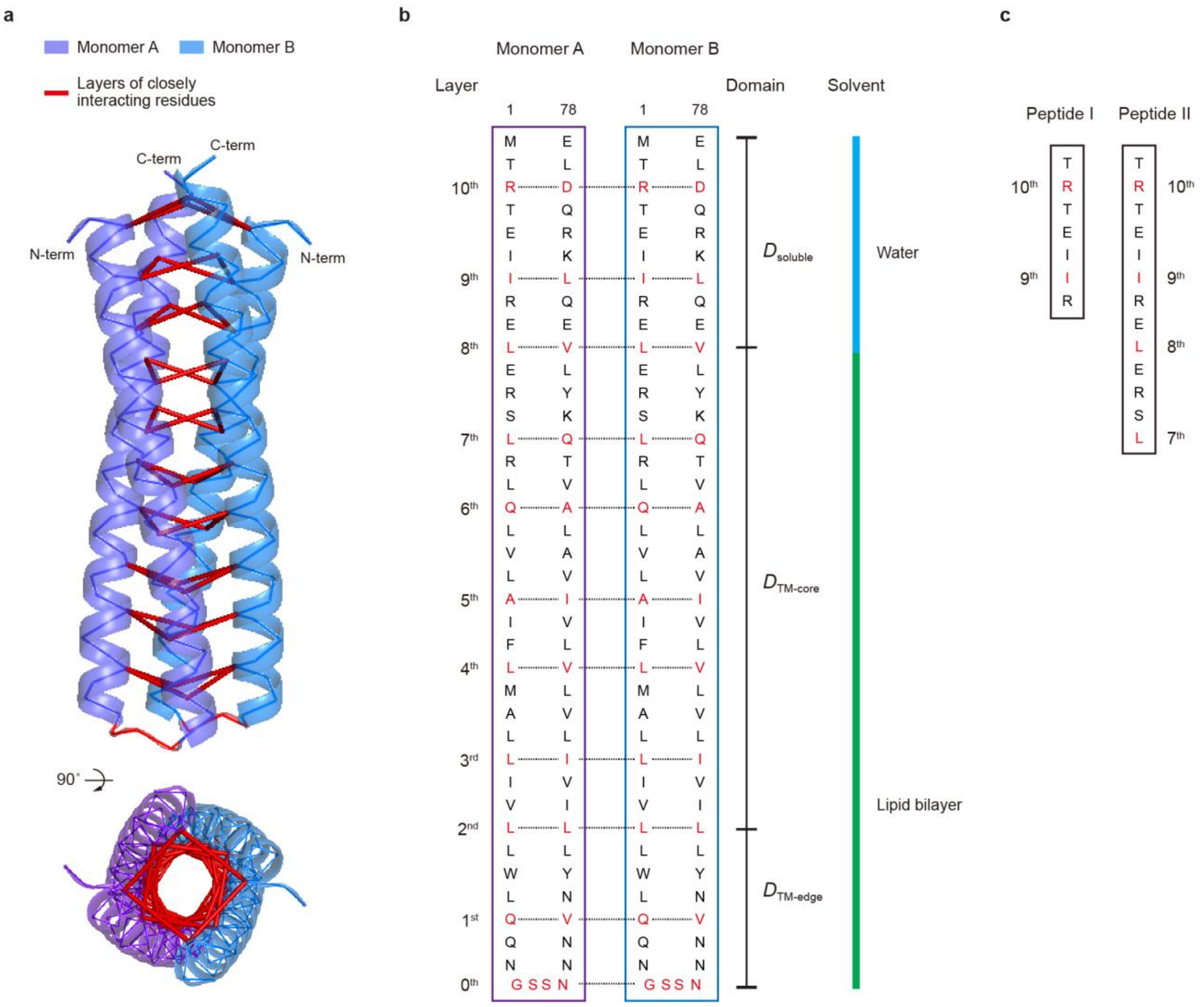


**Supplementary Fig. 4 | Protein interaction layers and peptides.** (**a**) Structure of the TMHC2 dimer formed by identical monomers showing the interaction layers (purple, monomer A; blue, monomer B). The red lines indicate the closely interacting inner residues at the dimerization interface. Each group of the residues are designated as an interaction layer. (**b**) Amino acid sequence of TMHC2 monomers. The residues of each layer are highlighted in red. (**c**) Amino acid sequence of peptides used in this work.

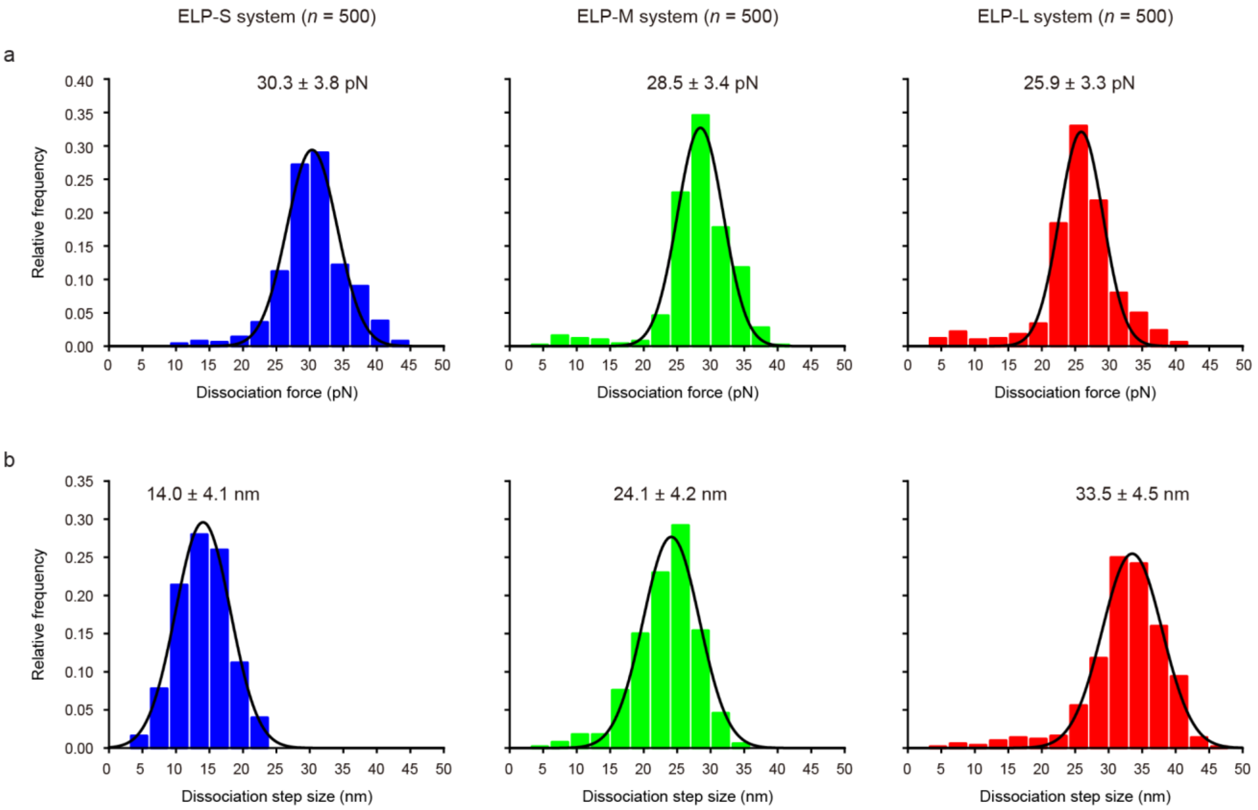


**Supplementary Fig. 5 | Dissociation forces and step sizes of TMHC2 dimer.** (**a**,**b**) Distributions of dissociation forces (**a**) and step sizes (**b**) in the force-cycle experiments with a constant magnet speed of 0.1 mm/s (each $n$ = 500 data points from 5 molecules). These data represent the complete dissociation from the $M$ to $D$ state. The distributions are fitted with Gaussian functions (peak ± SD).

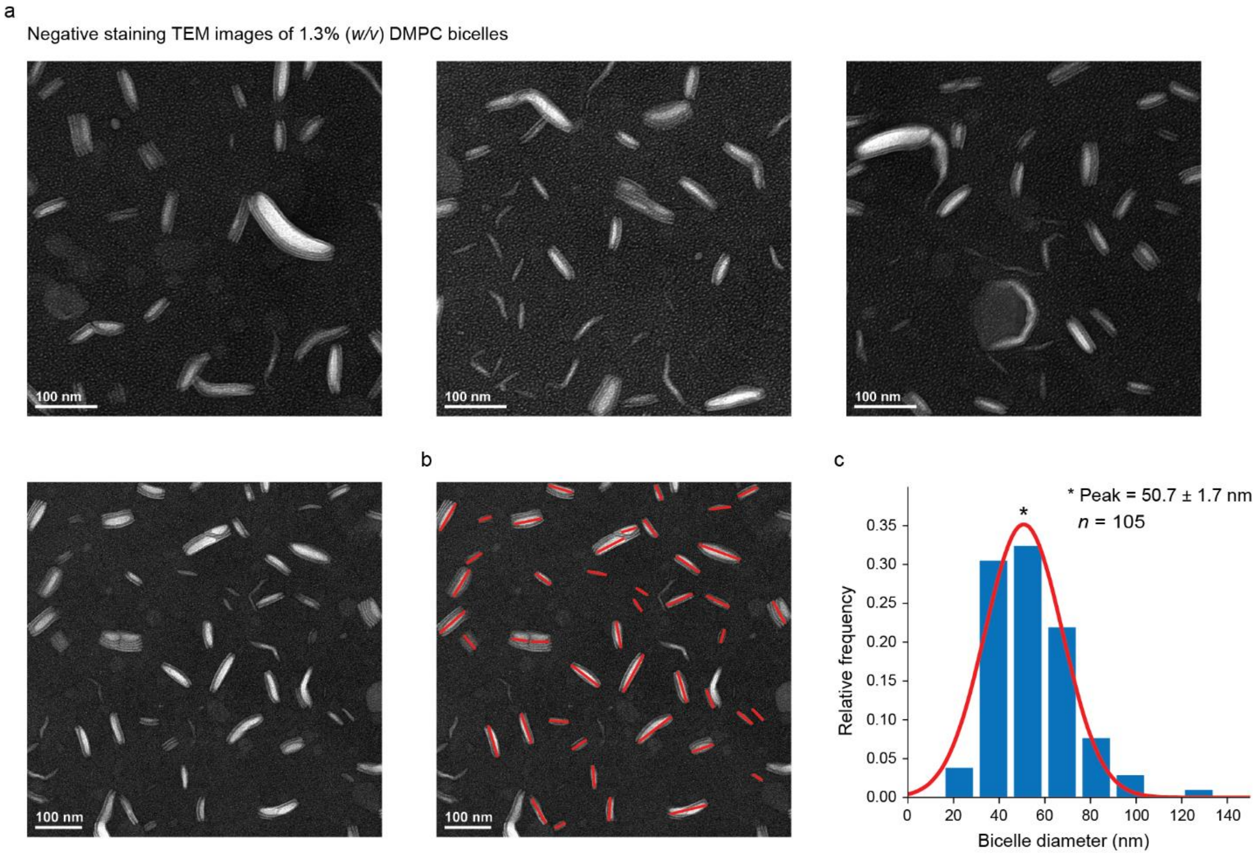


**Supplementary Fig. 6 | Bicelle size characterized using TEM.** (**a**) Negative staining transmission electron microscopy (TEM) images of 1.3% (w/v) DMPC bicelles. The scale bar indicates 100 nm. (**b**) Representative TEM image with red lines indicating the diameters of corresponding bicelles. (**c**) Size distribution of bicelles analyzed from the TEM images (*n* = 105 data points). The relative frequency of bicelle diameter values is fitted with a Gaussian function. The peak value, 50.7 ± 1.7 (SE) nm, represents the most probable diameter of the bicelles used in the single-molecule tweezer experiments.

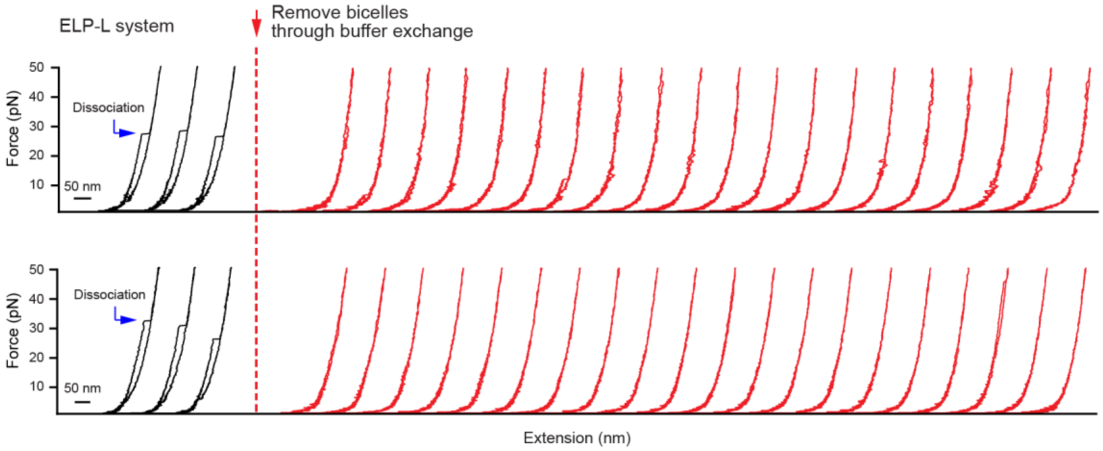


**Supplementary Fig. 7 | TMHC2 dimerization within lipid bilayers.** Force-extension curves of additional different molecules before and after the removal of bicelles (lipid bilayer discs). In the presence of bicelles (left, black traces), repetitive dissociation events of TMHC2 are observed over multiple cycles, indicating successful re-dimerization within bicelles. In contrast, the dissociation events of TMHC2 are not detected following the removal of bicelles through buffer exchange (right, red traces), indicating a failure of re-dimerization in the absence of bicelles. These results demonstrate that TMHC2 successfully dimerizes only with bicelles, underscoring the importance of reconstituting proper lipid bilayer environments.

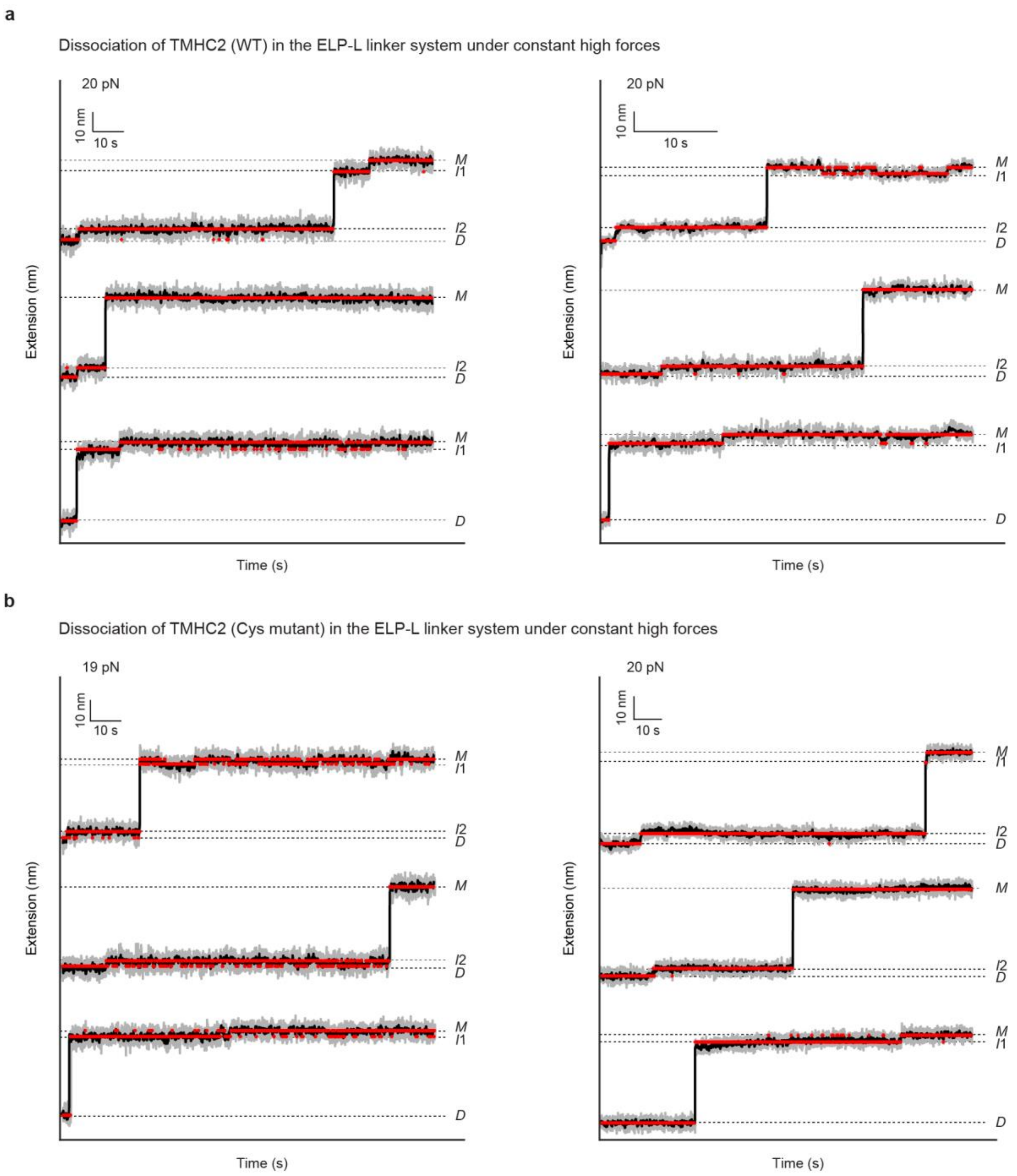


**Supplementary Fig. 8 | Representative high force traces in ELP-L system.** (**a**,**b**) Time-resolved extension traces at constant high forces for the wild-type (WT) of TMHC2 (**a**) and its Cys mutant (**b**). The gray and black traces represent the raw traces and the median-filtered traces with a 5-Hz window, respectively. The red traces represent the traces obtained using the hidden Markov modeling (HMM) with the optimal number of states. The optimal number of states was determined using the Bayesian information criterion (BIC) analysis.

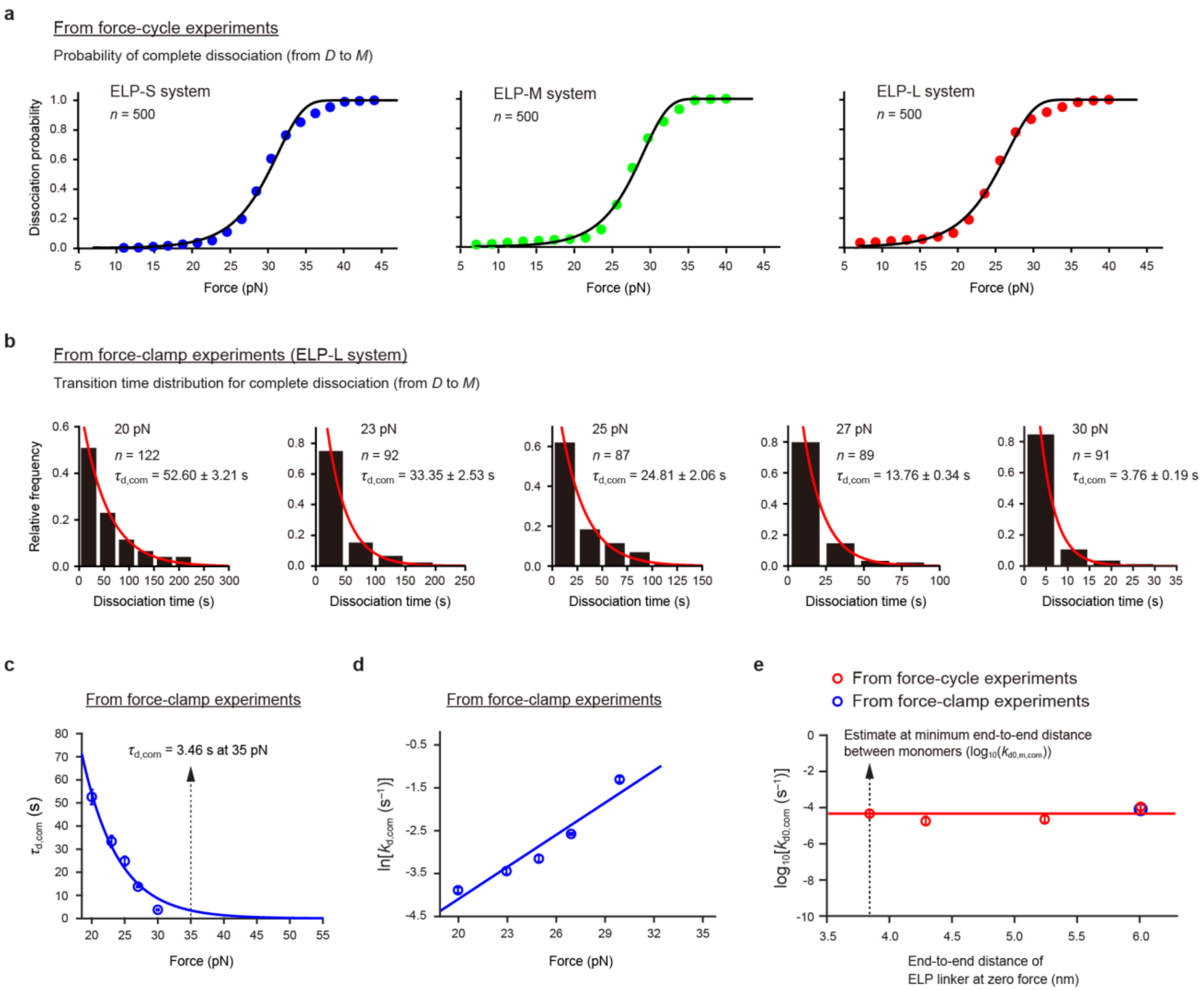


**Supplementary Fig. 9 | Transition kinetics for complete dissociation of TMHC2 dimer** (**a**) Probability of complete dissociation from the *D* to *M* state as a function of force for each ELP linker system (each $n$ = 500 traces from 5 molecules). These data were obtained from the force-cycle experiments. A regression analysis yields the dissociation rate constants at zero force, as shown in panel **e** (see Methods for details). (**b**) Distributions of complete dissociation times at constant forces. The complete dissociation times correspond to the total durations before transitioning from the *D* to *M* state at constant forces. These data were obtained from the force-clamp experiments. The number of data points is indicated in each panel ($n$ = 87–122 traces from 3 molecules). (**c**) Time constant for complete dissociation ($\tau_{\mathrm{d,com}}$) as a function of force. This data is obtained from data in panel **b**. The $\tau_{\mathrm{d,com}}$ value at 35 pN is estimated as 3.46 s, ensuring complete dissociation within 20 s. A 35 pN force was thus applied for 20 s to induce dissociation, followed by force reduction to characterize dimerization events in the low force-clamp experiments. (**d**) Rate constant for complete dissociation ($k_{\mathrm{d,com}}$) as a function of force. The $k_{\mathrm{d,com}}$ values, obtained as the reciprocals of $\tau_{d,com}$, are presented as their natural logarithm, *i.e.*, $\ln(k_{d,com})$. Extrapolation by a regression analysis yields the rate constant for complete dissociation at zero force ($k_{d0,com}$) (see Methods for details). (**e**) $\text{Log}_{10}(k_{d0,com})$ as a function of the end-to-end distance of the ELP linker at zero force. The data points from both the force-cycle and force-clamp experiments are presented. This data is used to estimate $k_{d0,com}$ at the minimum end-to-end distance between monomers, denoted as $k_{d0,m,com}$. The errors in panels **b**–**e** represent SE.

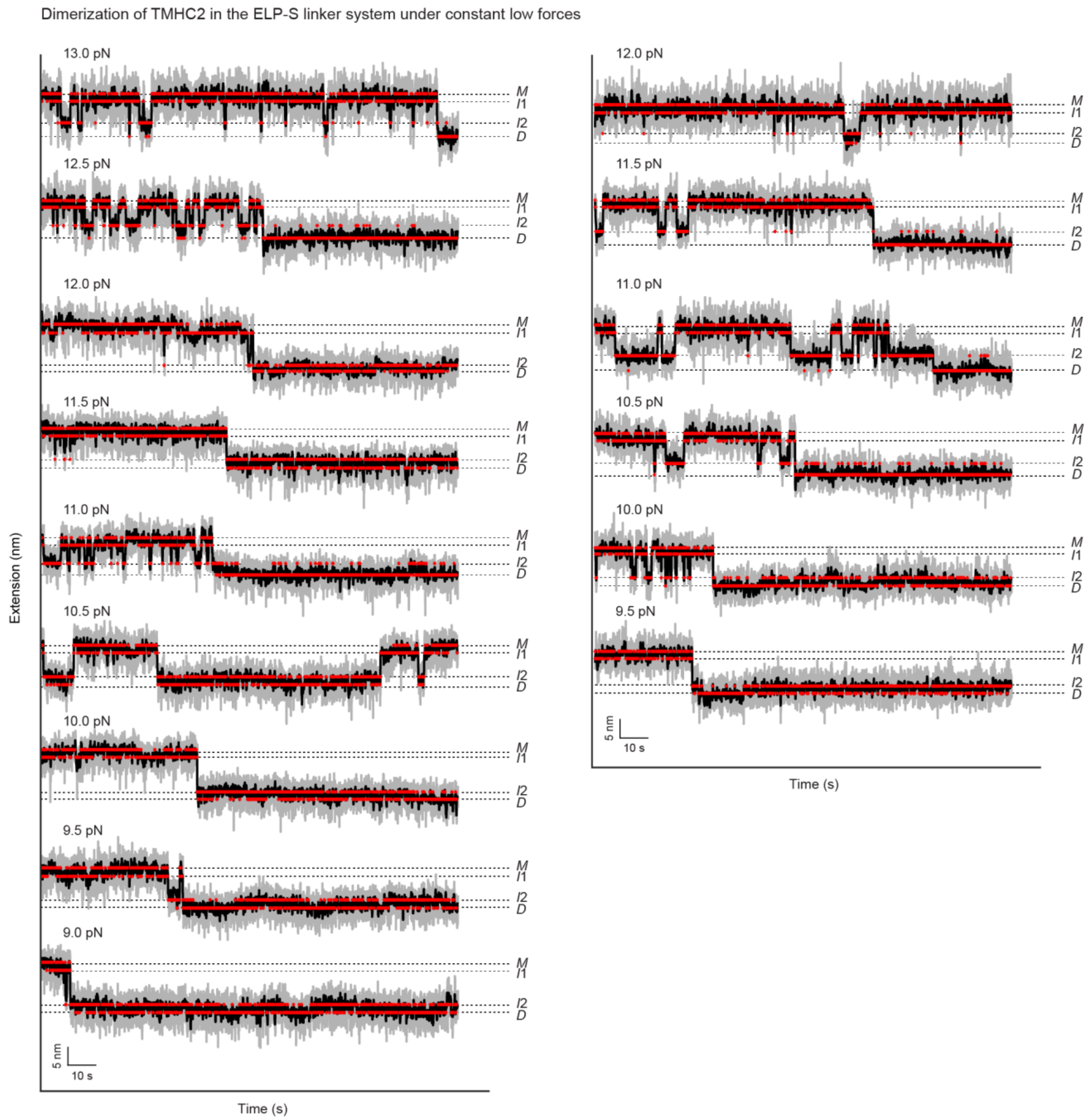


**Supplementary Fig. 10 | Representative low force traces in ELP-S system.** The gray and black traces represent the raw time-resolved extension traces and the median-filtered traces with a 5-Hz window at constant forces, respectively. The red traces represent the traces obtained using the hidden Markov modeling (HMM) with the optimal number of states. The optimal number of states was determined using the Bayesian information criterion (BIC) analysis.

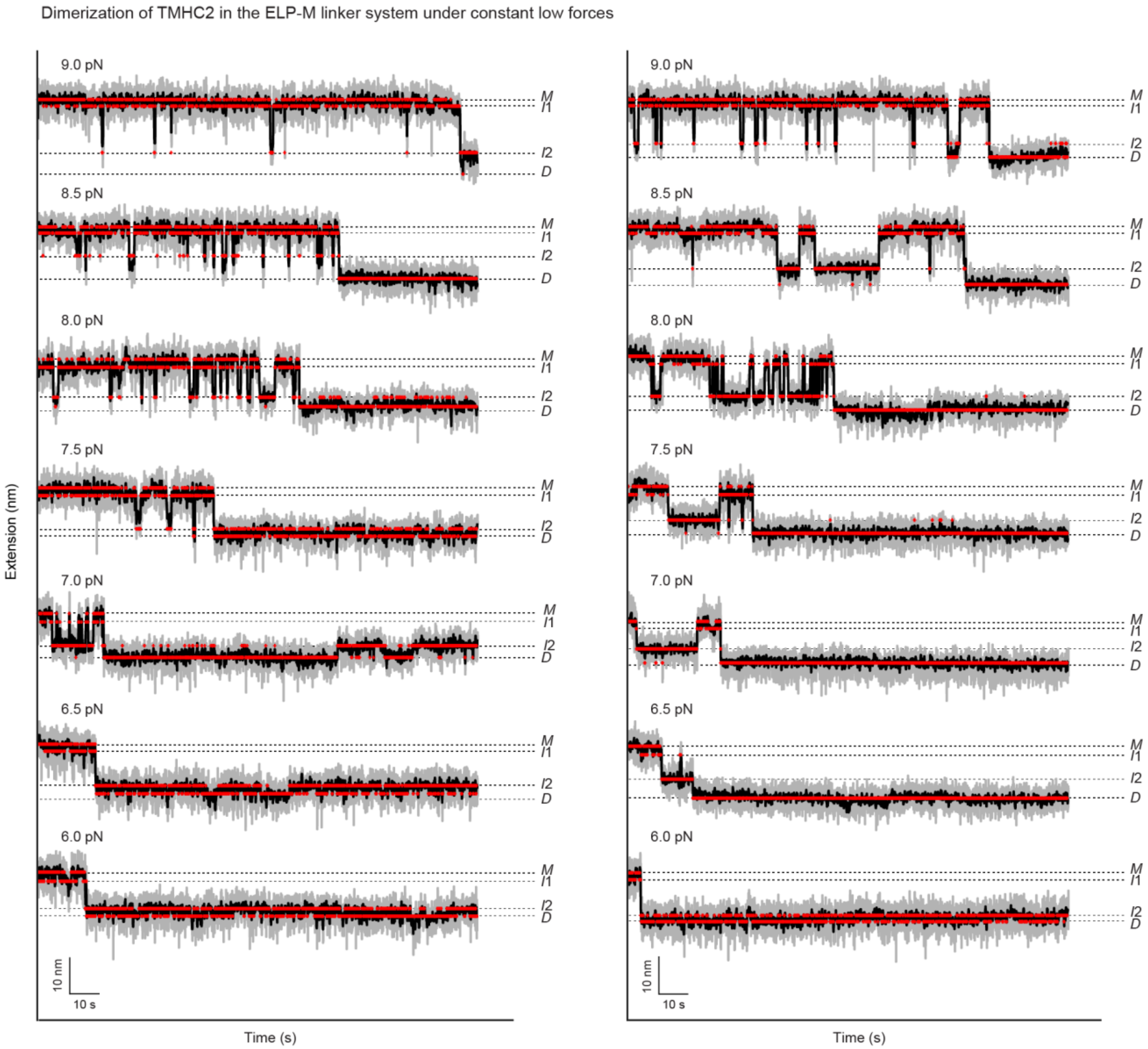


**Supplementary Fig. 11 | Representative low force traces in ELP-M system.** The gray and black traces represent the raw time-resolved extension traces and the median-filtered traces with a 5-Hz window at constant forces, respectively. The red traces represent the traces obtained using the hidden Markov modeling (HMM) with the optimal number of states. The optimal number of states was determined using the Bayesian information criterion (BIC) analysis.

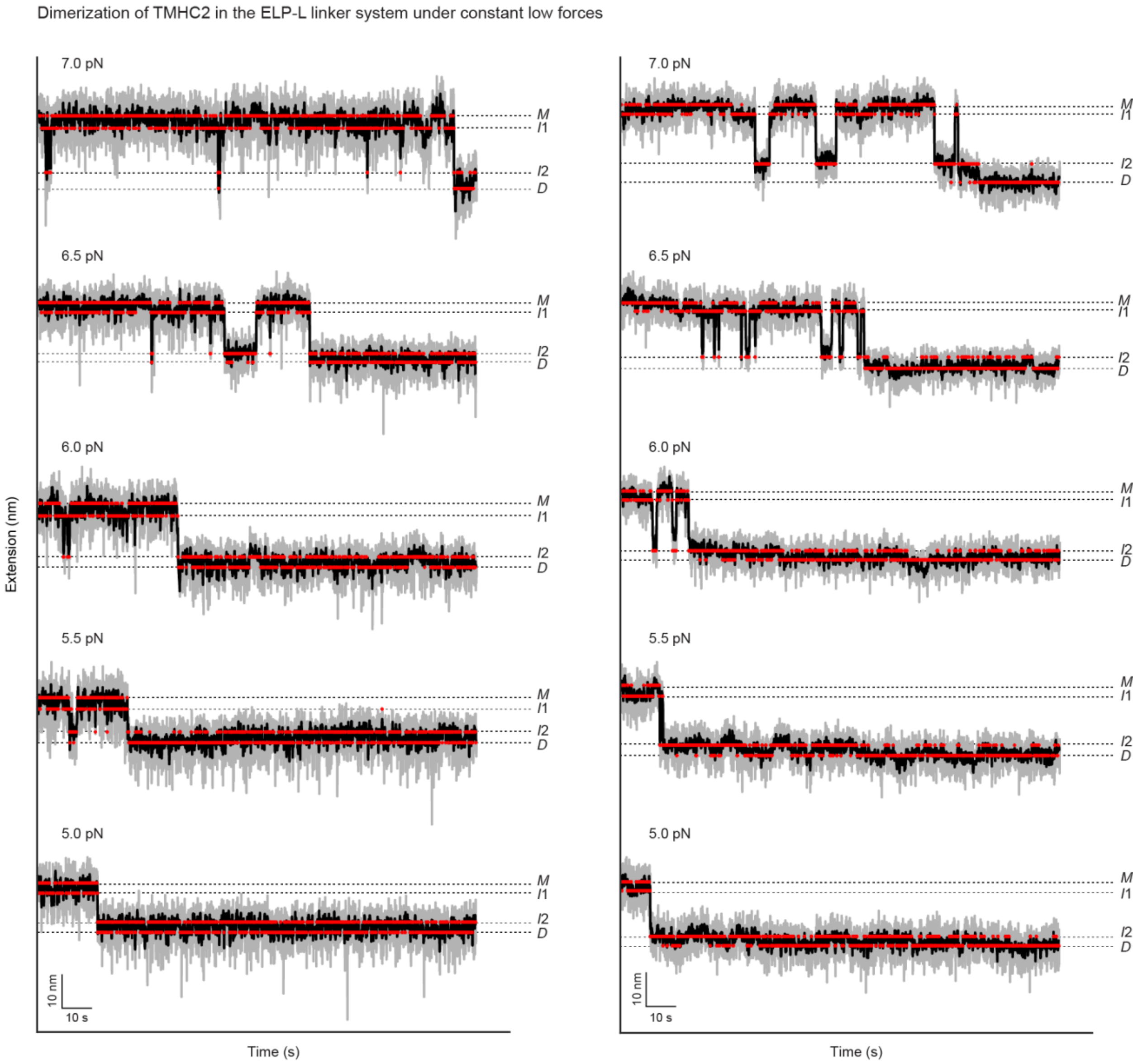


**Supplementary Fig. 12 | Representative low force traces in ELP-L system.** The gray and black traces represent the raw time-resolved extension traces and the median-filtered traces with a 5-Hz window at constant forces, respectively. The red traces represent the traces obtained using the hidden Markov modeling (HMM) with the optimal number of states. The optimal number of states was determined using the Bayesian information criterion (BIC) analysis.

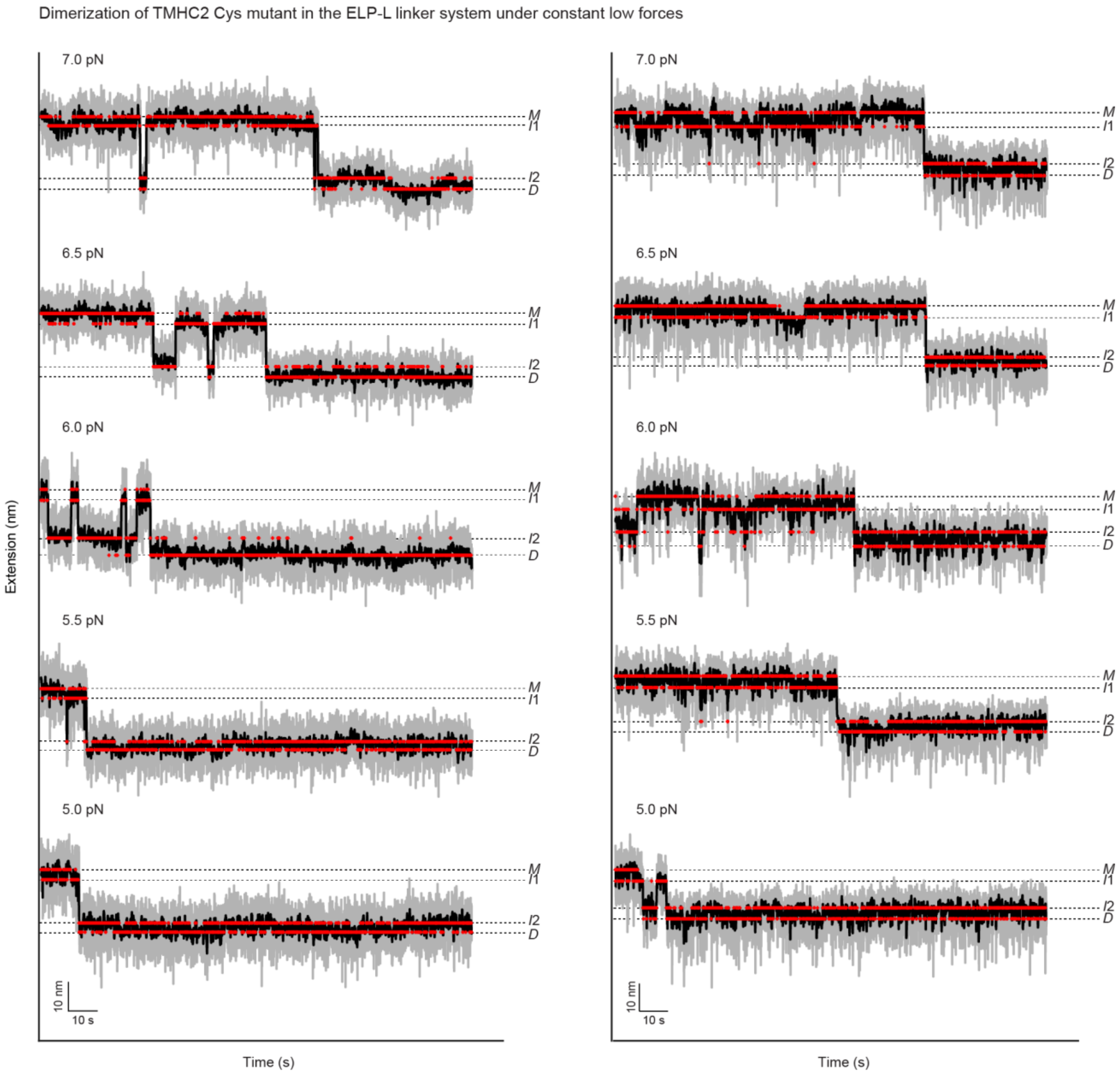


**Supplementary Fig. 13 | Representative low force traces of Cys mutant in ELP-L system.** The gray and black traces represent the raw time-resolved extension traces and the median-filtered traces with a 5-Hz window at constant forces, respectively. The red traces represent the traces obtained using the hidden Markov modeling (HMM) with the optimal number of states. The optimal number of states was determined using the Bayesian information criterion (BIC) analysis.

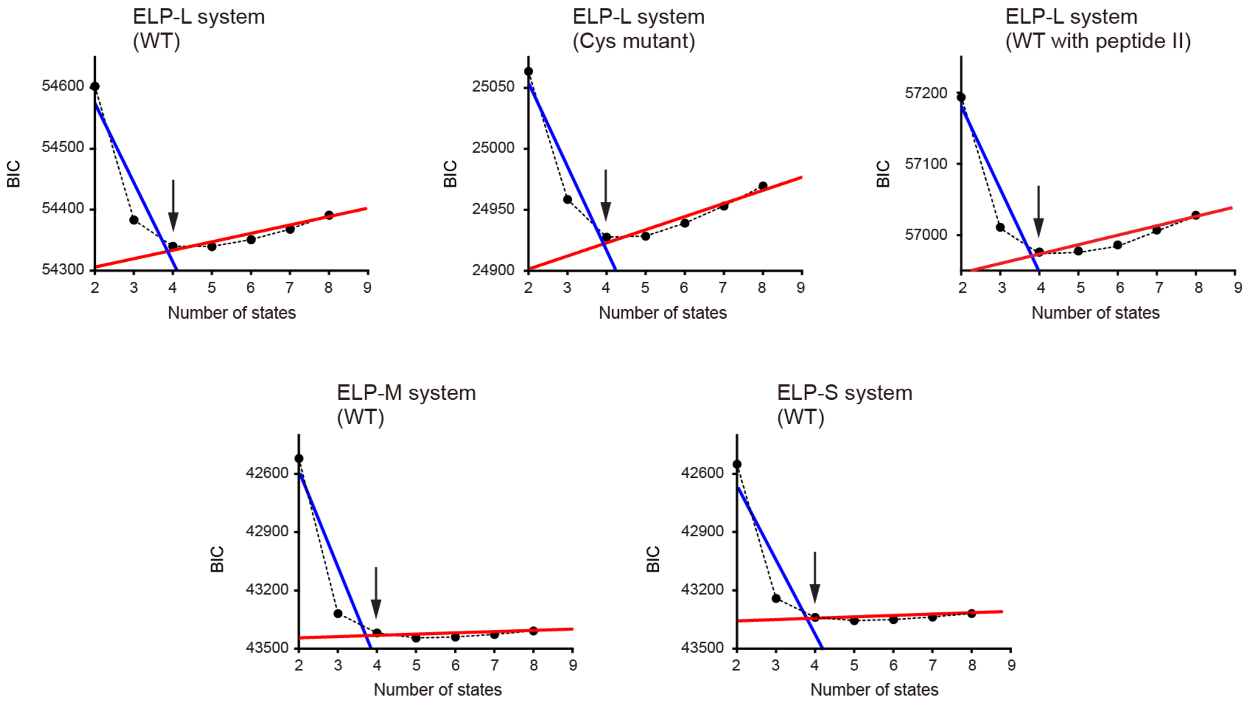


**Supplementary Fig. 14 | Bayesian information criterion (BIC) analysis.** The Bayesian information criterion (BIC) analysis was used to determine the optimal number of states. The BIC as a function of the number of states was obtained from the force-clamp extension traces showing the complete *M-D* transitions across all forces (*n* = 142 traces from 13 molecules for the ELP-L system (WT), *n* = 59 traces from 3 molecules for the ELP-L system (Cys mutant), *n* = 107 traces from 12 molecules for the ELP-L system (WT with peptide II), *n* = 150 traces from 8 molecules for the ELP-M system (WT), and *n* = 238 traces from 13 molecules for the ELP-S system (WT)). WT, Cys mutant, and WT with peptide II refer to the wild-type TMHC2, the Cys mutant of TMHC2, and the wild-type TMHC2 in the presence of peptide II, respectively. The optimal number of states was determined from the BIC plot as the elbow point, where the slope of the BIC changes substantially with the number of states (see Methods for details). The blue and red lines represent linear fits to visualize these significant slop changes, with arrows indicating the identified optimal number of states.

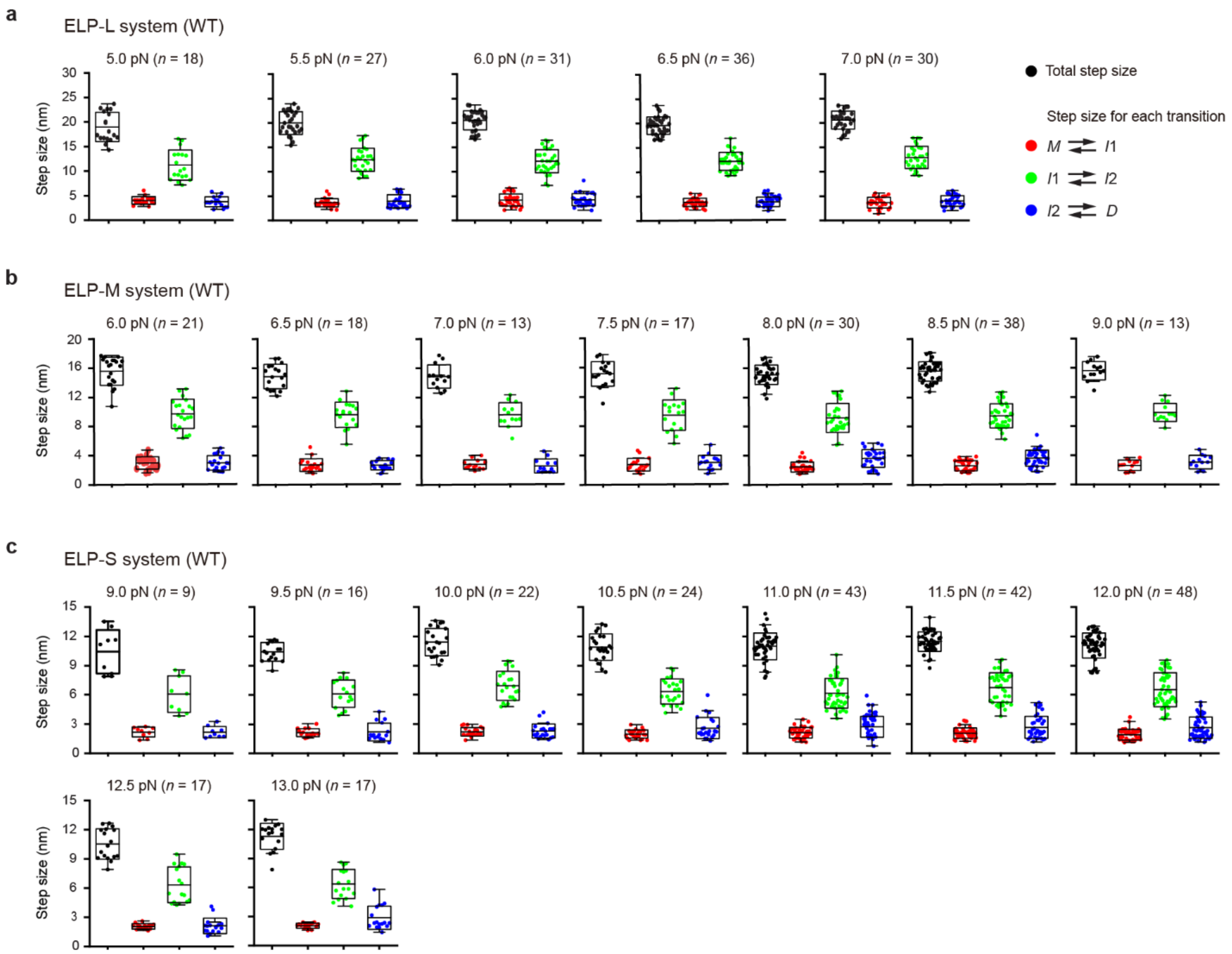


**Supplementary Fig. 15 | Transition step sizes of TMHC2.** (**a**–**c**) Step size distributions for the wild-type (WT) TMHC2 in the ELP-L (**a**), ELP-M (**b**), and ELP-S (**c**) linker systems. The data were obtained from the force-clamp extension traces showing the complete *M*-*D* transitions at each force. The number of data points is indicated in each panel ($n$ = 18–36 data points from 13 molecules for the ELP-L system, $n$ = 13–38 data points from 8 molecules for the ELP-M system, and $n$ = 9–48 data points from 13 molecules for the ELP-S system). The center lines of the boxes, the upper and lower edges of boxes, and the error bars represent the mean, standard deviation, and 1.5×interquartile range (IQR), respectively.

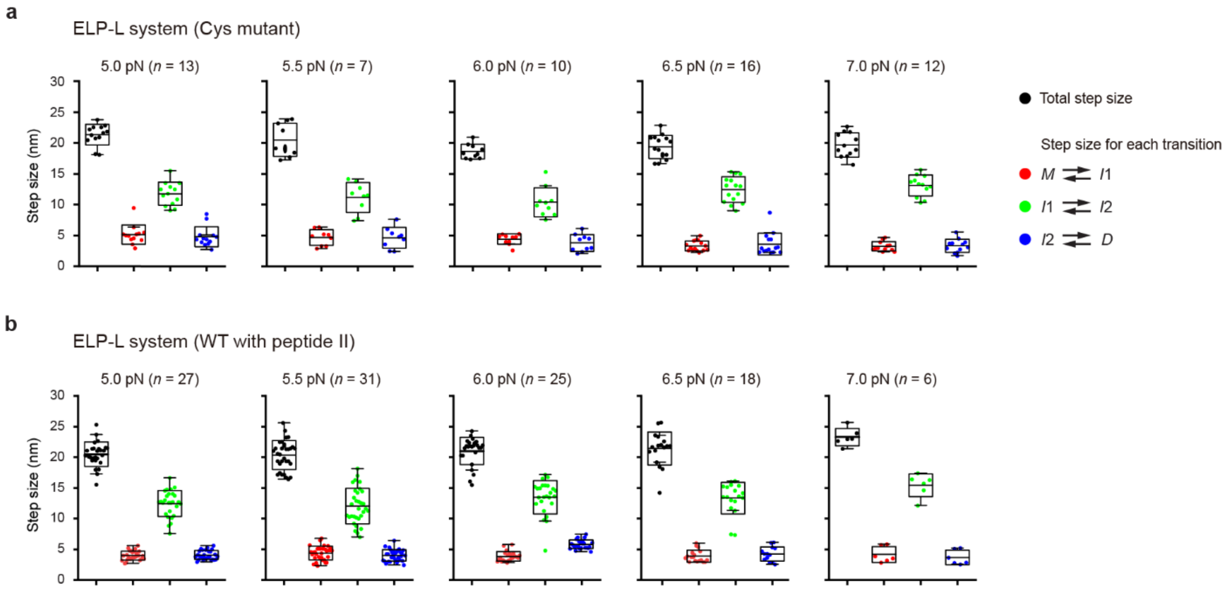


**Supplementary Fig. 16 | Transition step sizes of Cys mutant or WT with peptide II.** (**a,b**) Step size distributions for the TMHC2 Cys mutant (**a**) and the wild-type (WT) with peptide II (**b**) in the ELP-L linker system. The data were obtained from the force-clamp extension traces showing the complete *M*-*D* transitions at each force. The number of data points is indicated in each panel ($n$ = 7–16 data points from 3 molecules for the ELP-L system (Cys mutant) and $n$ = 6–31 data points from 12 molecules for the ELP-L system (WT with peptide II)). The center lines of the boxes, the upper and lower edges of boxes, and the error bars represent the mean, standard deviation, and 1.5×interquartile range (IQR), respectively.

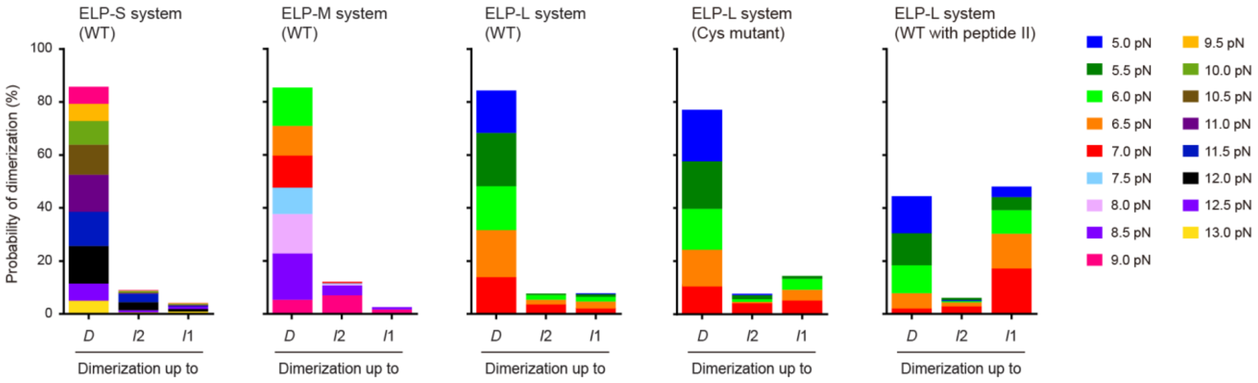


**Supplementary Fig. 17 | Probability of dimerization in force-clamp experiments.** The probability of dimerization up to each state was obtained from the force-clamp extension traces across all forces (*n* = 358 traces from 13 molecules for the ELP-S system (WT), *n* = 241 traces from 8 molecules for the ELP-M system (WT), *n* = 224 traces from 13 molecules for the ELP-L system (WT), *n* = 195 traces from 3 molecules for the ELP-L system (Cys mutant), and *n* = 342 traces from 12 molecules for the ELP-L system (WT with peptide II)). WT, Cys mutant, and WT with peptide II refer to the wild-type TMHC2, the Cys mutant of TMHC2, and the wild-type TMHC2 in the presence of peptide II, respectively.

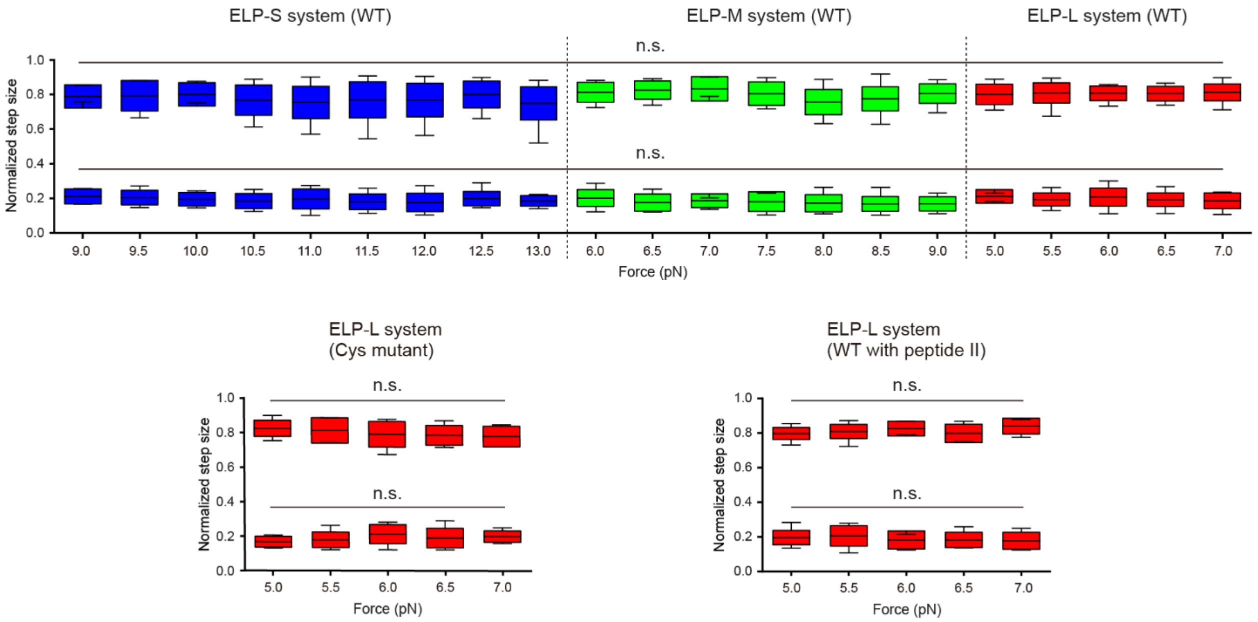


**Supplementary Fig. 18 | Normalized step sizes up to each intermediate state.** The step sizes for transitions from the *M* state up to the $I_1$ or $I_2$ state are normalized to the total step sizes for transitions from the *M* to *D* state. The step sizes were obtained from the force-clamp extension traces showing the complete *M*-*D* transitions at each force ($n$ = 9–48 data points from 13 molecules for the ELP-S system (WT), $n$ = 13–38 data points from 8 molecules for the ELP-M system (WT), $n$ = 18–36 data points from 13 molecules for the ELP-L system (WT), $n$ = 7–16 data points from 3 molecules for the ELP-L system (Cys mutant), and $n$ = 6–31 data points from 12 molecules for the ELP-L system (WT with peptide II)). WT, Cys mutant, and WT with peptide II refer to the wild-type TMHC2, the Cys mutant of TMHC2, and the wild-type TMHC2 in the presence of peptide II, respectively. The center lines of the boxes, the upper and lower edges of boxes, and the error bars represent the mean, standard deviation, and 1.5×interquartile range (IQR), respectively. One-way ANOVA with post-hoc Tukey test is used for step size comparisons (n.s. for $p > 0.05$).

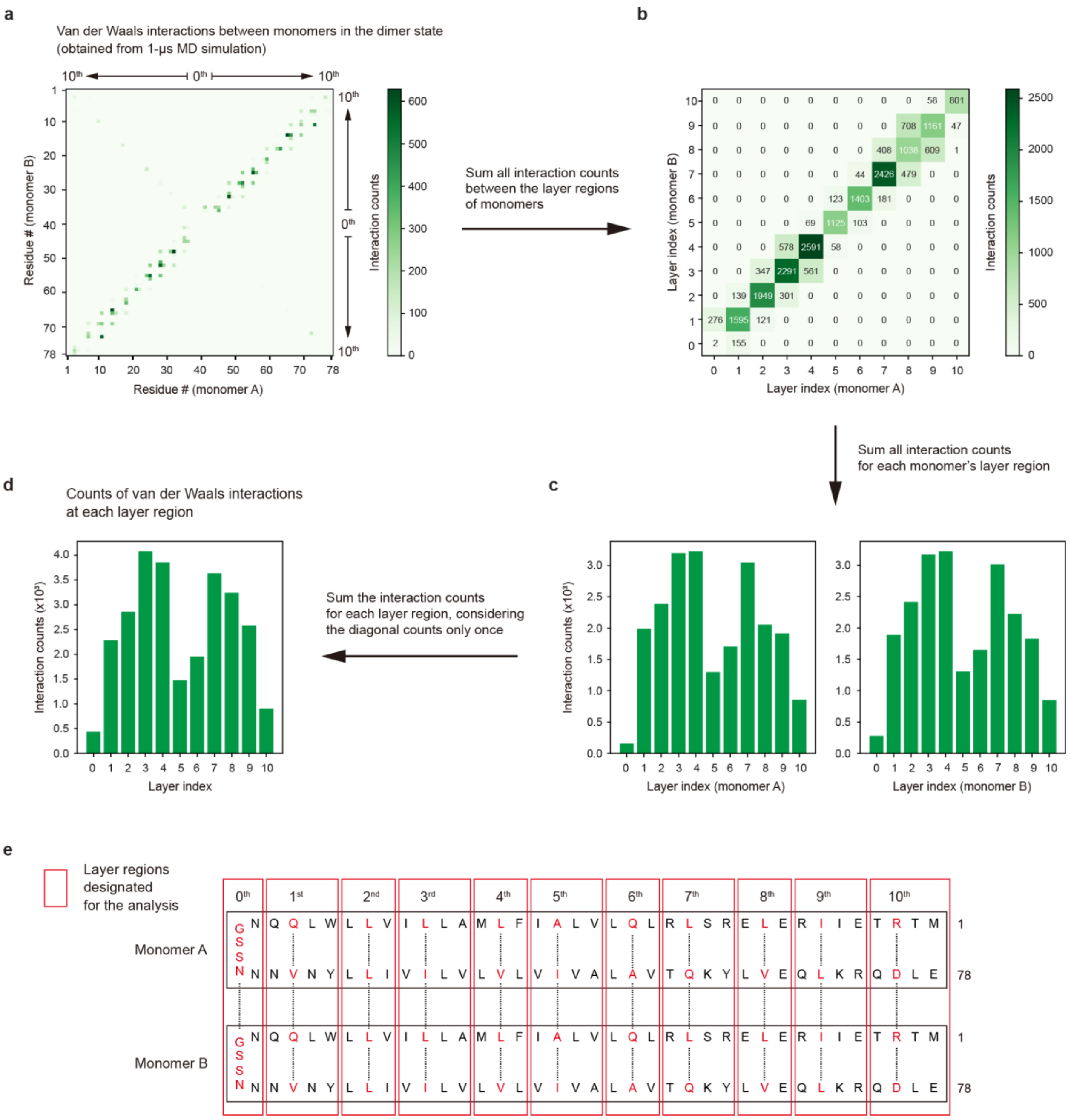


**Supplementary Fig. 19 | Analysis procedure of intermolecular interactions.** This figure presents the analysis procedure for intermolecular interactions between TMHC2 monomers shown in Fig. 3g, with van der Waals (vdW) interaction provided as an example case. Refer to the Methods section for details. (**a**) Heat map displaying the counts of vdW interactions between each residue of two monomers. (**b**) Heat map displaying the counts of vdW interactions between each interaction layer region of two monomers. The interaction layer regions are shown in panel **e**. (**c**) Bar plot displaying the counts of vdW interactions between one monomer's layer and all layers of the other monomer. The left plot shows the counts of interactions between each layer of monomer A and all layers of monomer B, while the right plot shows the counts of interactions between each layer of monomer B and all layers of monomer A. (**d**) Bar plot displaying the counts of vdW interactions at each layer region of both monomers. (**e**) Interaction layer regions designated for the analysis. The layer regions include residues within the red boxes. The closely interacting inner residues at the dimerization interface, highlighted in red, are located at the center position of each layer region, or the immediate next position.

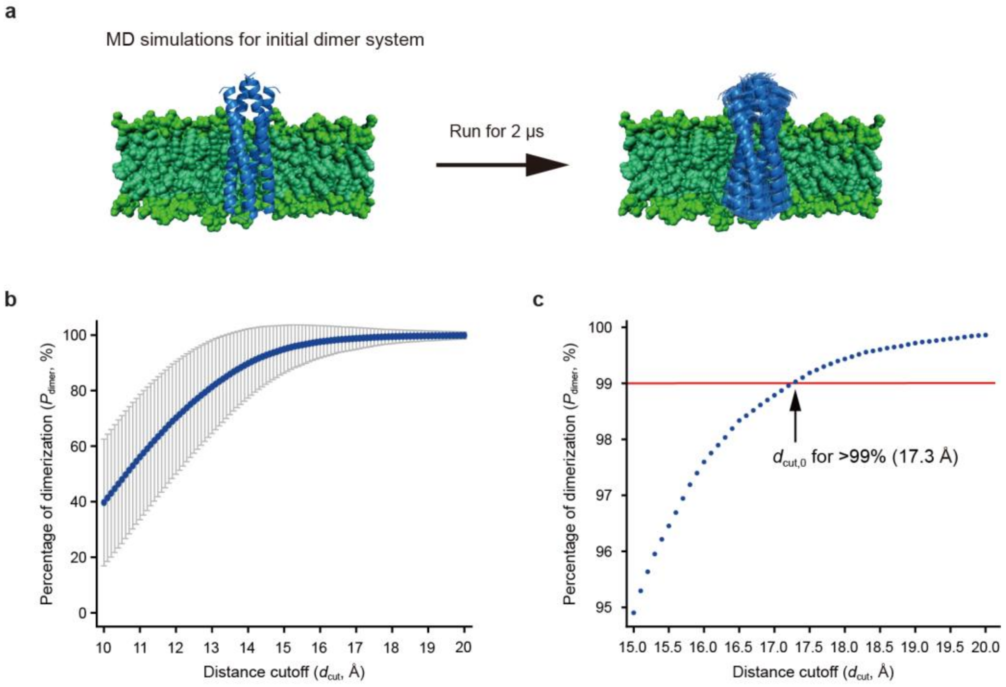


**Supplementary Fig. 20 | Determination of distance cutoff for binding at interaction layers.** (**a**) Coarse-grained molecular dynamics (CGMD) simulations for determining the distance cutoff for binding at the interaction layers ($d_{\mathrm{cut},0}$). (**b**) Percentage of dimerization ($P_{\mathrm{dimer}}$) for the initial dimer system as a function of arbitrary distance cutoff ($d_{\mathrm{cut}}$). For each simulation trajectory, the backbone bead coordinates of each layer's residues in two monomers are extracted at 1-ns intervals and averaged to obtain their center coordinates. The distance between the center coordinates in each layer ($d_{mm}$) is then calculated. The proportion of all layer's $d_{mm}$ values below an arbitrary distance cutoff $d_{\mathrm{cut}}$ ($P_{\mathrm{dimer}}$) is plotted as a function of $d_{\mathrm{cut}}$ ($n$ = 5 simulation trajectories; mean ± SD). (**c**) Close-up view around $P_{\mathrm{dimer}}$ = 99%. The $d_{\mathrm{cut}}$ at $P_{\mathrm{dimer}}$ = 99% is selected as the distance cutoff for binding at the interaction layers ($d_{\mathrm{cut},0}$; 17.3 Å).

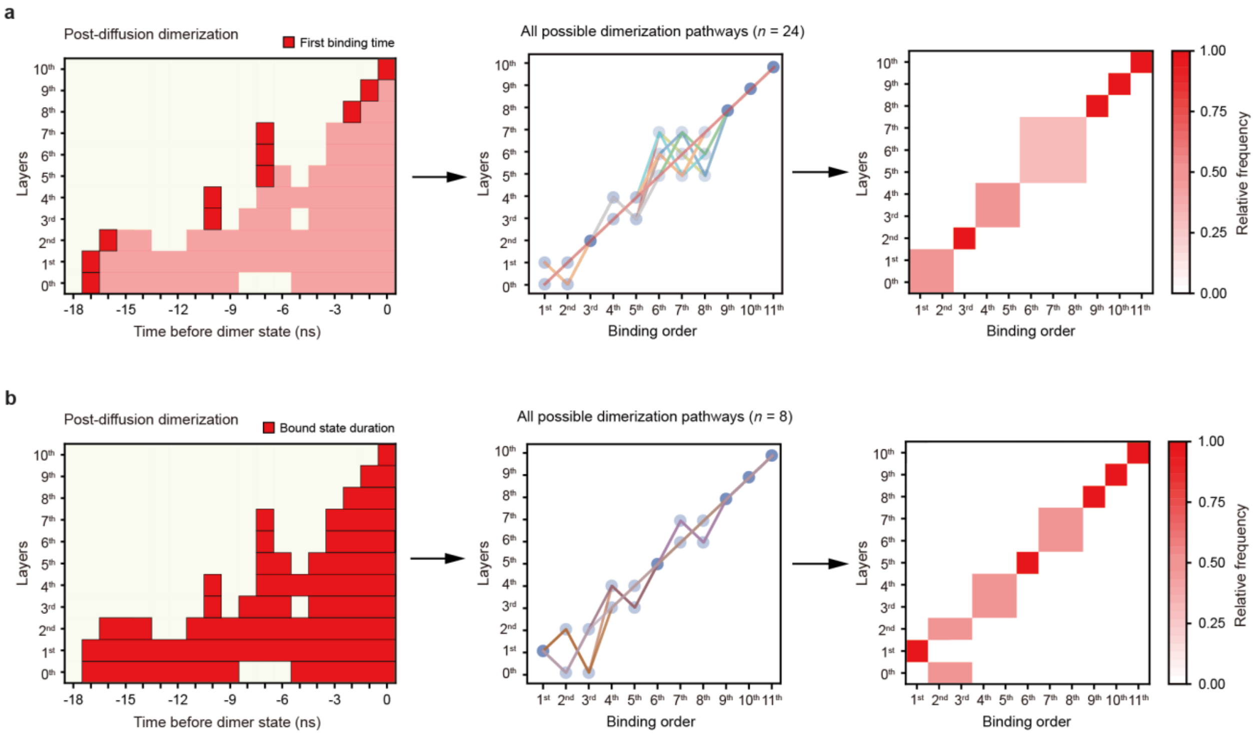


**Supplementary Fig. 21 | Analysis of binding order across interaction layers.** The binding order across the interaction layers during the post-diffusion dimerization period is analyzed using two different criteria: (**a**) the first binding time and (**b**) the total bound state duration (left). With each criterion, all possible binding pathways across the layers are identified (middle). The binding order across the layers is then quantified as the relative frequency of the rank order at each layer over all 80 trajectories that exhibited dimerization events (right). See Methods for details.

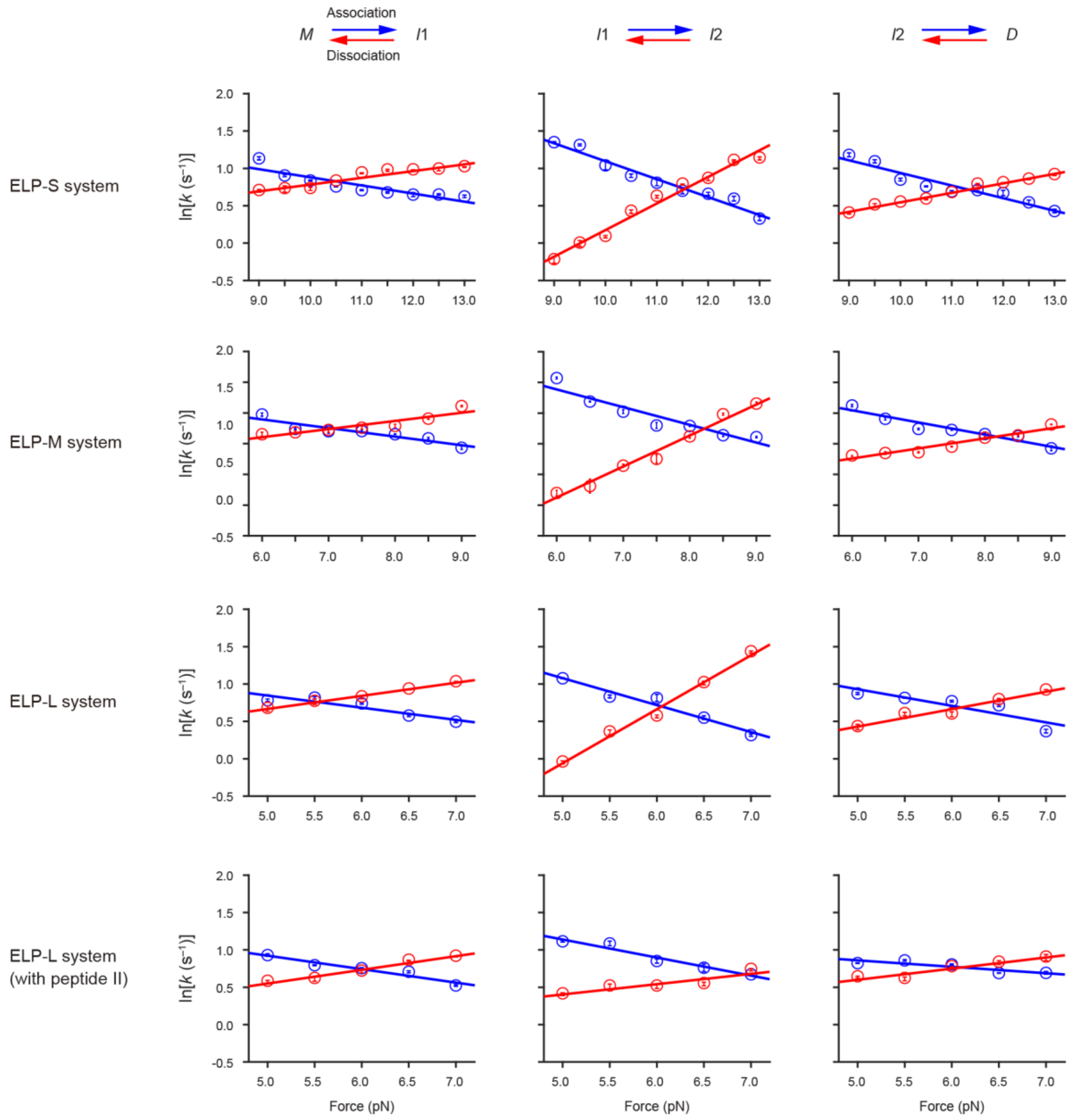


**Supplementary Fig. 22 | Force-dependent transition kinetics.** The rate constants for transitions between states ($k$) are presented as their natural logarithm ($\ln(k)$) as a function of force. The data for association and dissociation transitions are represented in blue and red, respectively. The regression analyses using the Bell-Evans model yield the rate constants for transitions at zero force ($k_0$). The error bars represent SE. See Methods for details.

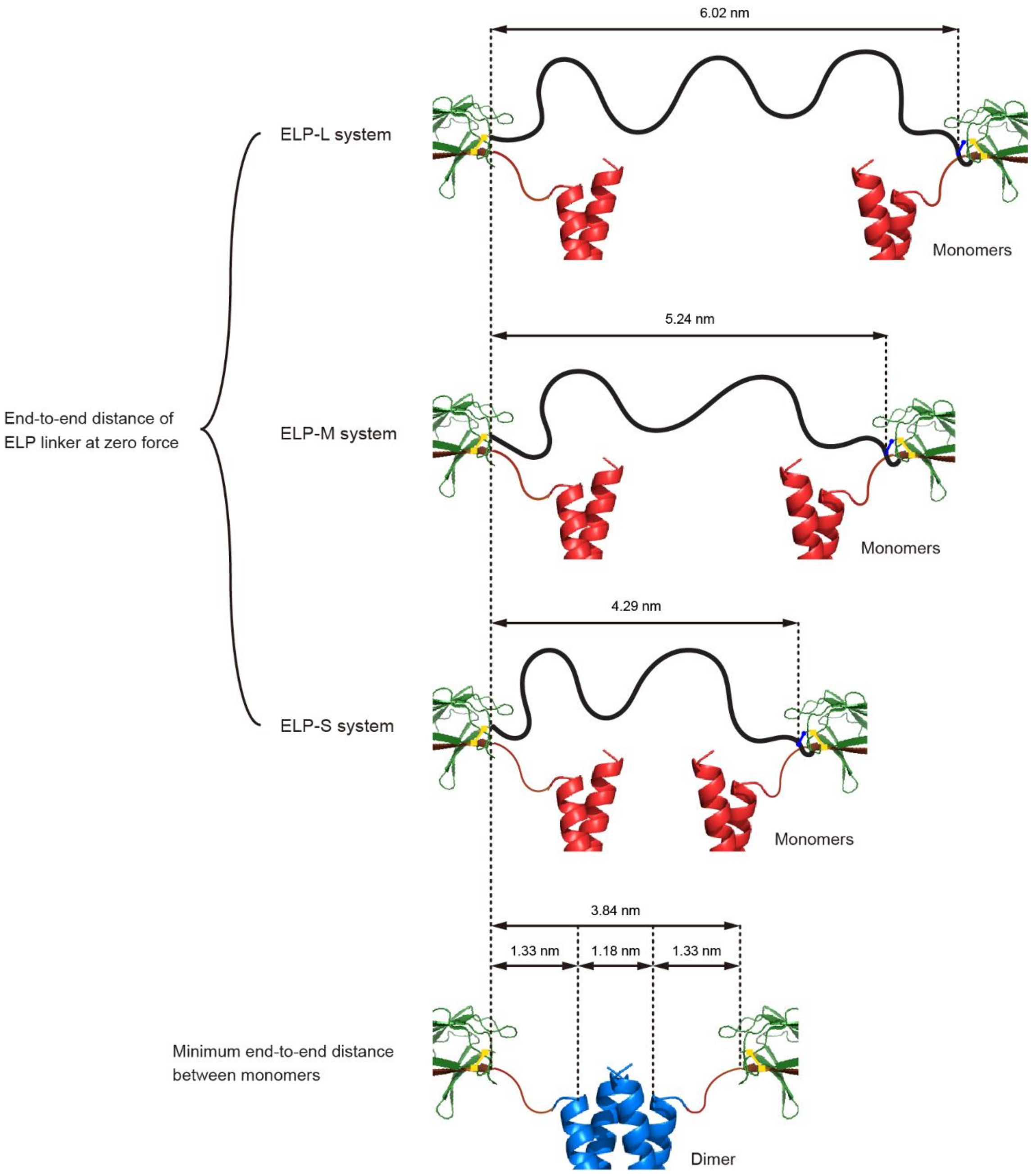


**Supplementary Fig. 23 | ELP linker extension at zero force and minimum monomer distance.** The end-to-end distances (extensions) of the ELP linkers at zero force ($d_{\text{elp-l}}$, $d_{\text{elp-m}}$, and $d_{\text{elp-s}}$) are estimated from the root mean square end-to-end distance of the worm-like chain (WLC) polymer ($d_{\text{rmsd}}$) for the polypeptide regions. The minimum end-to-end distance between monomers ($d_{\text{min}}$) is estimated from the end-to-end distance of the structured region of the dimer and the $d_{\text{rmsd}}$ of the linker regions flanking the dimer structure (see Methods for details).

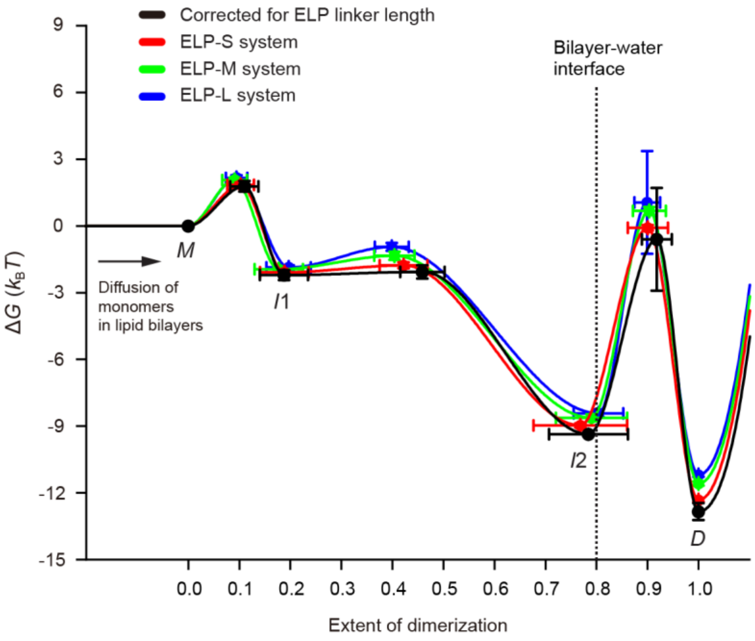


**Supplementary Fig. 24 | Reconstructed energy landscapes of TMHC2 dimerization.** The free energy landscape of TMHC2 dimerization in each ELP linker system, as well as the one corrected for the ELP linker length, are shown in different colors.

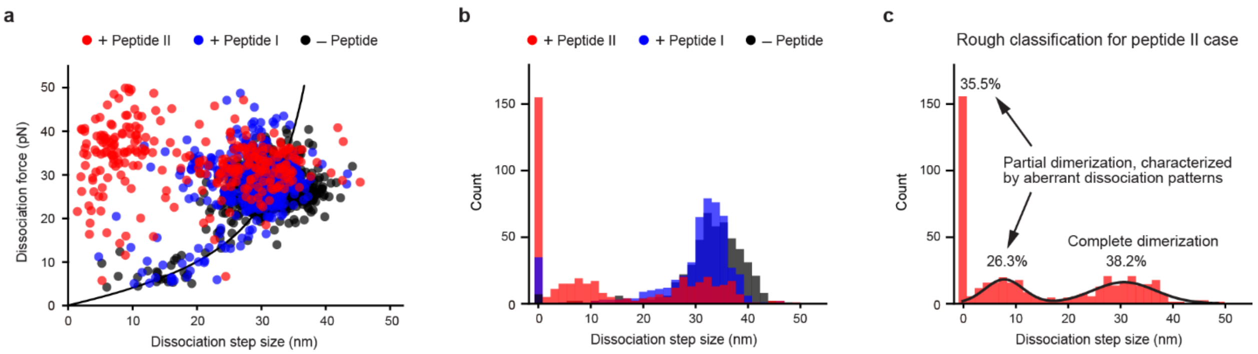


**Supplementary Fig. 25 | Dissociation forces and step sizes in the presence of peptides.** The data were obtained in the ELP-L linker system. (**a**) Scatter plot of dissociation forces and step sizes in the presence or absence of peptides during force-cycle experiments (*n* = 432–513 data points from 5–7 molecules). (**b**) Count histograms for dissociation step sizes (*n* = 432–513 data points from 5–7 molecules). (**c**) Rough classification of dimerization types for the peptide II case using the Gaussian mixture model (GMM) (*n* = 432 data points from 7 molecules). The populations on the leftmost and middle represent partial dimerization, characterized by aberrant dissociation patterns with either undetectable or small step sizes, while the population on the rightmost represents complete dimerization, consistent with the worm-like chain (WLC) model.

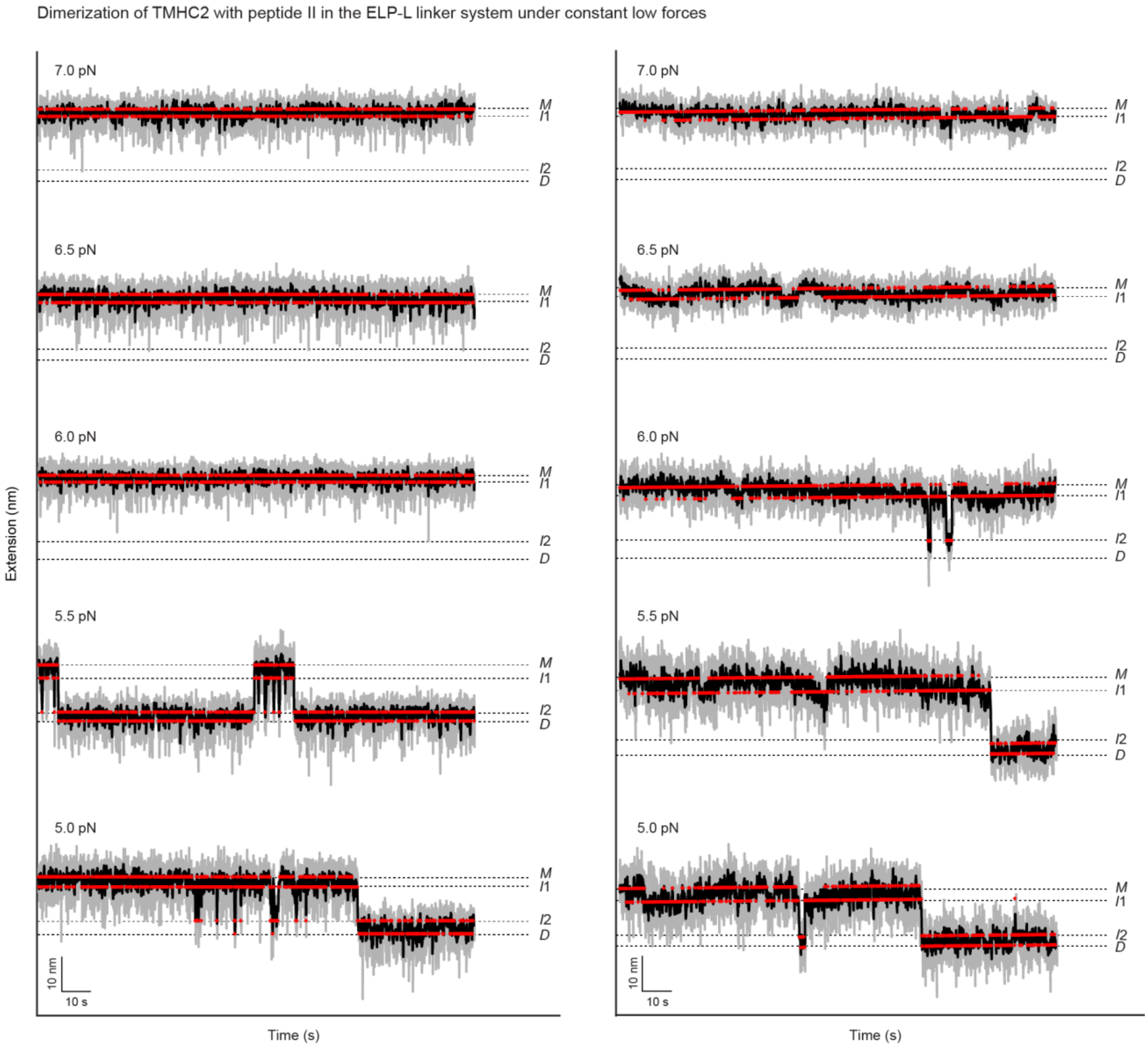


**Supplementary Fig. 26 | Representative low force traces in the presence of peptide II.** These traces were obtained in the ELP-L linker system. The gray and black traces represent the raw time-resolved extension traces and the median-filtered traces with a 5-Hz window at constant forces, respectively. The red traces represent the traces obtained using the hidden Markov modeling (HMM) with the optimal number of states. The optimal number of states was determined using the Bayesian information criterion (BIC) analysis.

**Supplementary Table 1 | Number of transitions in force-clamp experiments.** The numbers of association and dissociation transitions are indicated without and within parentheses, respectively. The numbers of traces and molecules are also shown on the right columns.

| Force (pN) | # of transitions | | | | | | # of traces | # of molecules |
|---|---|---|---|---|---|---|---|---|
| | Neighboring | | | Non-neighboring | | | | |
| | *M*–*I*$_1$ | *I*$_1$–*I*$_2$ | *I*$_2$–*D* | *M*–*I*$_2$ | *M*–*D* | *I*$_1$–*D* | | |
| ELP-S system | | | | | | | | |
| 13.0 | 1581 (1580) | 317 (305) | 664 (656) | 0 (0) | 0 (0) | 0 (0) | 24 | 9 |
| 12.5 | 1556 (1555) | 315 (302) | 1571 (1557 | 4 (5) | 0 (0) | 0 (0) | 30 | 9 |
| 12.0 | 3859 (3869) | 993 (967) | 2819 (2787) | 13 (3) | 0 (0) | 2 (0) | 65 | 13 |
| 11.5 | 2956 (2347) | 792 (761) | 2424 (2402) | 1 (0) | 0 (0) | 1 (0) | 60 | 13 |
| 11.0 | 2288 (2295) | 426 (392) | 2793 (2763) | 2 (0) | 0 (0) | 1 (0) | 52 | 13 |
| 10.5 | 1691 (1691) | 443 (414) | 3341 (3372) | 1 (1) | 0 (0) | 0 (0) | 42 | 13 |
| 10.0 | 669 (667) | 223 (202) | 2440 (2420) | 0 (0) | 0 (0) | 0 (0) | 34 | 13 |
| 9.5 | 892 (891) | 122 (109) | 1934 (1924) | 2 (0) | 0 (0) | 0 (0) | 26 | 13 |
| 9.0 | 553 (552) | 126 (120) | 1512 (1500) | 0 (0) | 0 (0) | 0 (0) | 25 | 13 |
| ELP-M system | | | | | | | | |
| 9.0 | 3778 (3788) | 407 (400) | 558 (552) | 0 (0) | 0 (0) | 0 (0) | 34 | 8 |
| 8.5 | 3134 (3130) | 508 (471) | 2444 (2415) | 0 (0) | 0 (0) | 0 (0) | 53 | 8 |
| 8.0 | 1131 (1125) | 305 (274) | 2059 (2033) | 0 (0) | 0 (0) | 0 (0) | 37 | 8 |
| 7.5 | 590 (588) | 96 (77) | 1503 (1494) | 0 (0) | 0 (0) | 0 (0) | 25 | 8 |
| 7.0 | 223 (220) | 90 (74) | 2353 (2343) | 0 (0) | 0 (0) | 0 (0) | 30 | 8 |
| 6.5 | 469 (466) | 137 (117) | 2151 (2140) | 0 (0) | 0 (0) | 0 (0) | 27 | 8 |
| 6.0 | 635 (631) | 244 (225) | 2014 (2010) | 0 (0) | 0 (0) | 0 (0) | 35 | 8 |
| ELP-L system | | | | | | | | |
| 7.0 | 2458 (2453) | 121 (98) | 1496 (1487) | 0 (0) | 0 (0) | 0 (0) | 44 | 13 |
| 6.5 | 3324 (3319 | 234 (197) | 2939 (2919) | 0 (0) | 0 (0) | 0 (0) | 50 | 13 |
| 6.0 | 1732 (1732) | 340 (312) | 3293 (3278) | 0 (0) | 0 (0) | 0 (0) | 45 | 13 |
| 5.5 | 979 (977) | 254 (224) | 4676 (4656) | 0 (0) | 0 (0) | 0 (0) | 48 | 13 |
| 5.0 | 847 (846) | 235 (211) | 4149 (4131) | 0 (0) | 0 (0) | 0 (0) | 37 | 13 |
| ELP-L system with peptide II | | | | | | | | |
| 7.0 | 8813 (8846) | 187 (178) | 599 (591) | 1 (0) | 0 (0) | 0 (0) | 77 | 12 |
| 6.5 | 7148 (7164) | 318 (303) | 1217 (1207) | 0 (0) | 0 (0) | 0 (0) | 71 | 12 |
| 6.0 | 5878 (5890) | 424 (396) | 4036 (4013) | 0 (0) | 0 (0) | 0 (0) | 69 | 12 |
| 5.5 | 4529 (4539) | 811 (784) | 5380 (5359) | 0 (0) | 0 (0) | 2 (0) | 62 | 12 |
| 5.0 | 2912 (2923) | 387 (369) | 4243 (4229) | 0 (0) | 0 (0) | 0 (0) | 63 | 12 |

**Supplementary Table 2 | Kinetic and energetic estimates at zero force.** The values were obtained from data in the force-clamp experiments.

| | | Transition | $k_0$ ($s^{-1}$) | $\Delta x^{\dagger}$ (nm) | $\Delta G^{\dagger}$ ($k_BT$) |
|---|---|---|---|---|---|
| ELP-S system | Association | $M$–$I_1$ | 7.16 ± 0.52 | 0.44 ± 0.11 | 1.84 ± 0.11 |
| | | $I_1$–$I_2$ | 33.0 ± 2.30 | 0.97 ± 0.10 | 0.31 ± 0.10 |
| | | $I_2$–$D$ | 13.8 ± 0.99 | 0.68 ± 0.11 | 8.89 ± 0.07 |
| | Dissociation | $D$–$I_2$ | 0.49 ± 0.01 | 0.51 ± 0.04 | 12.23 ± 0.02 |
| | | $I_2$–$I_1$ | 0.034 ± 0.003 | 1.44 ± 0.11 | 7.19 ± 0.11 |
| | | $I_1$–$M$ | 0.89 ± 0.04 | 0.37 ± 0.06 | 3.93 ± 0.09 |
| ELP-M system | Association | $M$–$I_1$ | 5.72 ± 0.41 | 0.57 ± 0.11 | 2.07 ± 0.11 |
| | | $I_1$–$I_2$ | 24.6 ± 3.2 | 1.22 ± 0.20 | 0.61 ± 0.15 |
| | | $I_2$–$D$ | 9.14 ± 0.71 | 0.79 ± 0.11 | 9.30 ± 0.08 |
| | Dissociation | $D$–$I_2$ | 0.48 ± 0.03 | 0.67 ± 0.09 | 12.3 ± 0.06 |
| | | $I_2$–$I_1$ | 0.031 ± 0.003 | 2.08 ± 0.14 | 7.28 ± 0.12 |
| | | $I_1$–$M$ | 0.81 ± 0.06 | 0.54 ± 0.01 | 4.02 ± 0.11 |
| ELP-L system | Association | $M$–$I_1$ | 5.26 ± 0.55 | 0.66 ± 0.12 | 2.15 ± 0.13 |
| | | $I_1$–$I_2$ | 17.8 ± 2.2 | 1.46 ± 0.14 | 0.93 ± 0.16 |
| | | $I_2$–$D$ | 7.69 ± 0.04 | 0.90 ± 0.22 | 9.470 ± 0.005 |
| | Dissociation | $D$–$I_2$ | 0.48 ± 0.04 | 0.94 ± 0.10 | 12.2 ± 0.09 |
| | | $I_2$–$I_1$ | 0.026 ± 0.003 | 2.92 ± 0.14 | 7.48 ± 0.14 |
| | | $I_1$–$M$ | 0.81 ± 0.02 | 0.71 ± 0.03 | 4.02 ± 0.08 |
| ELP-L system with peptide II | Association | $M$–$I_1$ | 6.19 ± 0.48 | 0.73 ± 0.09 | 1.99 ± 0.11 |
| | | $I_1$–$I_2$ | 10.5 ± 0.92 | 0.98 ± 0.10 | 1.46 ± 0.12 |
| | | $I_2$–$D$ | 3.63 ± 0.27 | 0.35 ± 0.09 | 10.2 ± 0.08 |
| | Dissociation | $D$–$I_2$ | 0.86 ± 0.06 | 0.60 ± 0.09 | 11.7 ± 0.07 |
| | | $I_2$–$I_1$ | 0.75 ± 0.07 | 0.56 ± 0.12 | 4.09 ± 0.13 |
| | | $I_1$–$M$ | 0.69 ± 0.04 | 0.74 ± 0.06 | 4.18 ± 0.10 |
| ELP linker length corrected | Association | $M$–$I_1$ | 7.53 ± 1.7 | 0.39 ± 0.05 | 1.78 ± 0.24 |
| | | $I_1$–$I_2$ | 36.9 ± 0.25 | 0.84 ± 0.09 | 0.14 ± 0.19 |
| | | $I_2$–$D$ | 15.0 ± 5.68 | 0.62 ± 0.04 | 8.76 ± 0.40 |
| | Dissociation | $D$–$I_2$ | 0.48 ± 0.002 | 0.38 ± 0.33 | 12.2 ± 0.003 |
| | | $I_2$–$I_1$ | 0.031 ± 0.002 | 1.01 ±0.76 | 7.29 ± 0.08 |
| | | $I_1$–$M$ | 0.82 ± 0.02 | 0.27 ± 0.04 | 3.99 ± 0.03 |
